# A splice-switching antisense oligonucleotide approach for pediatric genetic epilepsies

**DOI:** 10.1101/2025.10.22.683934

**Authors:** Haley B. Dame, Hema Kopalle, Belen Miñana, Zin Klaft, Jaclyn B. Fahey, David C. McWatters, Vama Rao, Rushil Suresh, Stefan Aigner, Michael F. Hammer, Christopher J. Yuskaitis, Megan Wong, Nicole Teaney, Crystal Zhang, Mike Thompson, Ben Lehner, C. Frank Bennett, Juan Valcárcel, Gene W. Yeo, Christopher B. Burge, Chris G. Dulla, Madeleine J. Oudin

## Abstract

Variants in ion channel genes are common causes of pediatric epilepsy, often leading to intractable seizures, developmental delay and other comorbidities, which increases risk of death. Pathogenic variants in the *SCN8A* gene, which encodes a voltage-gated sodium channel critical for action potential generation in the brain, account for ∼1% of genetic epilepsies. The voltage sensor in *SCN8A* domain 1 is encoded by one of two developmentally-regulated mutually exclusive alternative exons, 5N and 5A. We observe that variants in these exons are more likely to cause infantile spasms, a severe seizure type, than variants elsewhere in *SCN8A*, and that some pathogenic variants affect exon 5 splicing, impacting patient phenotype. Molecular and evolutionary analyses implicate the exon sequences of these and other voltage-gated ion channel alternative exons in splicing regulation. We identified antisense oligonucleotides (ASOs) that shift splicing of *SCN8A* exon 5N to 5A or vice versa. These ASOs normalize neuronal activity in patient-derived iPSC neurons, and reduce seizures and motor impairment and extend lifespan in a new exon 5N mutant mouse model. Our results demonstrate that splice-switching ASOs can effectively reduce the expression of pathogenic isoforms and rescue both seizure and non-seizure phenotypes. Similar approaches should be applicable to pediatric genetic epilepsies caused by mutations in other ion channel alternative exons.

## Introduction

Epilepsy is one of the most common neurological disorders, affecting approximately 1% of people^1^. Pathogenic variants in the four main voltage-gated sodium channels (VGSCs) expressed in the brain (*SCN1A*, *SCN2A*, *SCN3A*, and *SCN8A*) are leading causes of developmental epileptic encephalopathies (DEE), with those in *SCN8A* accounting for up to 1% of all diagnoses^2^. *SCN8A* encodes the alpha subunit of the abundant VGSC Na_v_1.6, which localizes to the axon initial segment in neurons and is necessary for the regulation and generation of action potentials^3–5^. Variants with gain-of-function (GOF) properties in *SCN8A* – which are typically de novo – increase Na_v_1.6 activity, leading to intractable epilepsy, developmental delays and intellectual disability. Further, 90% of patients exhibit hypotonia and motor impairment, contributing to multiple comorbidities that affect peripheral organs such as the lungs, the gut or bones^6,7^. Death by respiratory failure is in fact more common than seizure-related deaths in DEE patients with intractable epilepsy and profound physical disabilities^8–11^. Published mouse models carrying *SCN8A* GOF variants exhibit seizures and reduced lifespan, but do not always replicate other comorbidities such as motor impairment^12–14^. Current treatments for *SCN8A* epilepsy include drugs that block VGSC channels or enhance GABAergic signaling. However, these drugs have limited efficacy and primarily target seizure reduction rather than developmental delay or comorbidities, emphasizing the need for new treatments for *SCN8A* epilepsy^15^.

Voltage-gated ion channels contribute to cardiac disease, epilepsies, ataxias and cancer^16^ and are often regulated by the alternative splicing of mutually exclusive exons (MXEs). In the CNS, the *SCN1A*, *SCN2A*, *SCN3A*, and *SCN8A* genes undergo conserved, developmentally regulated alternative splicing of MXEs 5N and 5A. These exons encode distinct but similar versions of the voltage sensor in domain I. Modeling suggests that the domain I voltage sensor is the rate-limiting step for release of the inactivation domain of many VGSCs, highlighting its functional importance^17^. In *SCN8A*, early developmental inclusion of the 5N “neonatal” exon shifts gradually to predominant inclusion of the 5A “adult” exon during childhood, and a similar shift occurs in mouse *Scn8a*^18^. In rodent ND7/23 cells, the Na_v_1.6A isoform exhibited a higher peak current, hyperpolarized voltage-dependence of activation, and a mildly hyperpolarized voltage-dependence of inactivation compared to Na_v_1.6N^19^, all consistent with modestly increased function. Whether developmental regulation of ion channel MXE splicing impacts brain development, whether its dysregulation impacts patient outcomes, how this splicing is regulated, and whether this can be leveraged therapeutically have not been investigated.

Splice-switching antisense oligonucleotides (ASOs) can be used to precisely alter splicing and/or mRNA abundance to impact protein function, and ASO therapies have been approved for various neuromuscular disorders^20–23^. Neurodevelopmental disorders are an attractive target for ASO development due to the efficient uptake of ASOs by neurons throughout the brain following injection into the cerebrospinal fluid^24^. Here, we present a splice-switching ASO approach for pediatric genetic epilepsies with variants in MXE exons. We show that pathogenic variants in *SCN8A* can alter the splicing of exon 5, which impacts clinical outcomes. We developed ASOs that induce a splicing switch from exon 5N to 5A (or vice versa) and can correct Na_v_1.6 function in iPSC-derived neurons from a patient with a pathogenic exon 5N mutation, S217P. Lastly, we developed a novel *Scn8a* exon 5N mouse model with the mutation S217P^+/-^, which has seizures, hypotonia, mis-regulated splicing, decreased levels of parvalbumin-positive (PV-INs), and reduced lifespan. These phenotypes are all improved with a splice-switching ASO. Together, our results highlight a novel disease-modifying approach to treat severe pediatric epilepsies which improves both seizure and non-seizure outcomes.

## Results

### Variants in SCN8A exon 5 lead to a severe DEE phenotype, with exon 5 variant location impacting seizure control

Exon 5 of *SCN8A* encodes a 31-amino acid portion of the Na_v_1.6 domain 1, covering part of segments 3 and 4, which is the voltage sensing domain of the protein (Fig. 1a). Developmentally regulated alternative splicing occurs between two mutually exclusive exons (MXEs), 5N and 5A. The two exons differ at 19 nucleotide positions but encode peptides that differ by only 2 amino acids (Fig. 1b). We identified a cohort of 52 patients diagnosed with variants in exon 5 of *SCN8A* (Table S1). Because both exons are identical in length, the cDNA position does not distinguish which alternative exon harbors the variant. From case studies and registry entries that report the chromosomal location, we identified the alternative exon location for 25 patients (Fig. 1c).

We compared the phenotypes of the exon 5 cohort to a general *SCN8A* patient group (*n*=240), from the International SCN8A registry ^25^. Overall, 91.7% of patients in the *SCN8A* cohort develop seizures^26^, compared to 98% in the exon 5 cohort (Fig. 1d). Median seizure onset is 3 months for patients with known GOF variants in *SCN8A*, but 18 months for patients with loss of function (LOF) variants^26,27^, with earlier seizure onset associated with more severe outcomes. Median seizure onset for patients with variants in exon 5 is 4 months, similar to typical GOF seizure onset, and several exon 5 patient variants studied exert a GOF effect on Na_v_1.6 activity^19,28,29^ (Fig. 1e). All patients with variants in exon 5 had some level of developmental delay, with 6 patients described with mild or moderate delays and intellectual disability, and 31 patients with global or severe developmental delay (Table S1). Patients with *SCN8A* variants develop a wide range of seizure types, including infantile spasms, which are known to lead to high mortality by age 5 with poor developmental outcomes and a high likelihood of developing other seizure types^8,30^. Strikingly, we observed a 3.5-fold increased incidence of infantile spasms in patients with variants in exon 5 compared to the rest of the gene (Fig. 1f).

We then compared a smaller cohort of patients with known variants in either exon 5N or 5A where seizure control status was reported. While median age of seizure onset was 5 months for both groups (Fig. 1g), patients with variants in exon 5N had a higher likelihood of seizure control compared to patients with variants in exon 5A (Fig. 1h). These data suggest that the developmental switch in splicing from exon 5N to 5A reduces pathogenic effects associated with 5N variants over time, yielding increased seizure control. Conversely, patients with pathogenic 5A variants may experience increasing disease severity as levels of 5A mRNA and protein increase with age. Despite reduced seizure burden, a majority of patients with 5N variants struggle with severe global developmental delays, suggesting that seizure reduction alone is not sufficient to rescue behavioral and cognitive phenotypes, reverse neurological impairments or restore normal physiological function.

**Figure 1:**
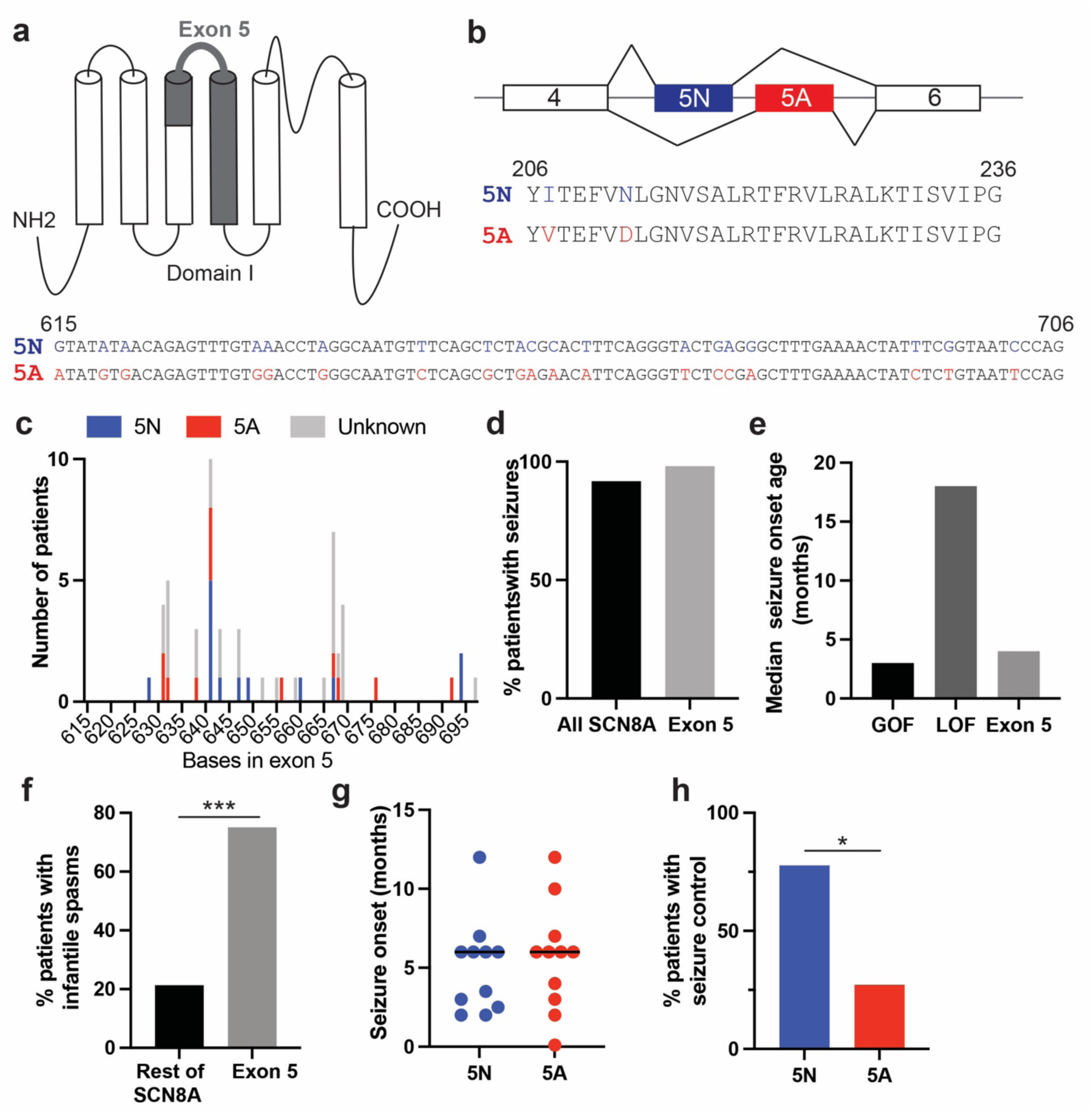
Pathogenic variants in exon 5 of SCN8A lead to severe phenotypes, but outcome is influenced by location of variant. **a**) Diagram of location of exon 5 (in dark grey) in Domain I of SCN8A **b**) DNA and amino acid sequence of exon 5N and 5A **c**) Known pathogenic patient variants, *n*=52 patients. Location of the variant in exons 5N or 5A, when known, is indicated by the color key. Comparing the entire SCN8A patient population to patients with exon 5 mutations in terms of presence of seizures (**d**), median age of seizure onset (**e**) and incidence of infantile spasms (**f**). 22 patients have known exon location for mutations and reported seizure control status: while age of seizure onset is the same in patients with 5N or 5A mutations (**g**), a significantly higher proportion of patients with variants in 5N have seizure control compared to patients with variants in 5A (**h**). Statistical significance by hypergeometric test: * p<0.05 and *** p<0.001.

### Sequence conservation and splicing regulation of mutually exclusive exons are tightly associated

While many of these variants have been characterized for their effects on Na_v_1.6 biophysical properties^19,28,29^, the effect of exon 5 variants on splicing remains unknown. Splicing regulation can be mediated by RNA elements located in exons or introns^31^. To obtain clues about splicing regulation for variant interpretation and ASO design, we examined nucleotide conservation in ion channel genes using alignments of 241 mammalian genomes^32^. Protein-coding constraint predominantly impacts the first and second positions of codons while the third site is often synonymous and therefore less constrained, a pattern ubiquitously observed in coding exons^33^. Most exons in *SCN8A* follow this pattern, with position 3 about half as conserved as 1 and 2, but in exons 5N and 5A conservation at third sites is far higher than for other exons, approaching that at positions 1 and 2 (Fig. 2a, 2c). This pattern suggests a role for these positions outside of protein coding, such as in regulation of splicing or expression. The *SCN8A* gene contains a second pair of MXEs, exons 18N/18A, of which 18A is coding and 18N contains a premature termination codon, yielding a nonfunctional isoform. We observe a similar pattern of high position 3 constraint in both 18N and 18A (Fig. 2b,2c). The high conservation of 18N in the absence of protein coding function further supports the presence of strong regulatory constraint on these MXEs.

To explore MXE conservation more broadly, we considered all MXE pairs present in human VGSCs and evolutionarily related voltage-gated Ca^2+^ channel (VGCC) genes^34^, which contain many MXE pairs (Fig. S1a). Strikingly, 12 of 14 Na^+^ channel MXEs and 16 of 16 Ca^2+^ channel MXEs have significantly higher constraint at third positions than other exons in the same gene (Fig. 2c shows the ratio of third site constraint between MXE exons and these other exons). The sole exceptions were *SCN1A* exons 5N and 5A. Notably, *SCN1A* exon 5N is not expressed – i.e. is a pseudoexon – in a number of mammalian species including mouse and other rodents^17,18^, explaining its low conservation at all 3 codon positions (Fig. S1d). *SCN1A* exon 5A is conserved and expressed across mammals, but is constitutively spliced in mammals that have lost exon 5N and shows conservation at positions 1 and 2 but less so at 3 (Fig. S1d). Thus, the excess conservation of third positions of codons is universally observed in VGSC and VGCC MXE pairs that undergo conserved mutually exclusive splicing but is reduced in cases where splicing is lost or becomes constitutive.

To further explore this association, we examined exon evolution in the Ca_v_1.X calcium channel family of *CACNA1-C, -D, -F and -S. CACNA1C* and *-D* each possess multiple pairs of MXEs, including 8a/8b and 31a/31b (Fig. S1b). *CACNA1F* and -*S* have each lost one exon 8 and one exon 31, retaining the other as a constitutive exon (Fig. S1b). These “singleton” or “widow” exons and their MXE counterparts are nearly identical in amino acid sequence (Fig. S1c), implying stable protein-level constraints. However, while the 8 *CACNA1C/D* MXEs have high third site constraint, the four singleton exons in *CACNA1F/*S have completely lost this conservation, resembling other constitutive exons found in these genes (Fig. 2c). Together, these observations establish that excess third site conservation is tightly associated with alternative splicing of MXEs in Na^+^ and Ca^+2^ channel genes, implicating these conserved bases in regulation of splicing of these exons.

To better understand exon 5N’s splicing regulation, we generated a mini-gene reporter of human *SCN8A* containing exons 2-6 (Fig. S2a) and conducted saturation mutagenesis to assess the impact on splicing. Exon 5N is identical between human and mouse, suggesting similar regulation. A library was built to assess the effects of every single-base substitution in exon 5N, as well as hundreds of double mutants (at flanking residues or at longer range) and all possible deletion mutants of 1, 3, 6 or 21 nt in length. The library was transfected into glioblastoma T98G cells and the single-exon inclusion product was isolated and deep sequenced. The relative inclusion level of exon 5N of each individual mutant was compared to that of the reference sequence by comparing the frequency of each variant in the spliced mRNA product including exon 5N, relative to its frequency at the DNA level in the original library. The results identify numerous positions at which substitutions impact splicing, with a majority of substitutions reducing 5N inclusion, suggesting the presence of several exonic splicing enhancer (ESE) elements in the exon (Fig. 2d, Table S2). Further, the majority of deletions that impacted exon 5N splicing decreased its inclusion, particularly for deletions involving the central region of the exon, further supporting the presence of ESEs (Fig. 2d).

Plotting the known variants from our cohort of exon 5N or exon 5 (with 5A versus 5N status unknown) patients, we found that several variants increased 5N inclusion in the minigene, while others had no splicing impact or decreased 5N inclusion (Fig. 2e). Considering that most pathogenic *SCN8A* mutations are GOF in nature, increased inclusion of the exon containing the variant is likely to exacerbate disease severity by increasing the levels of mutant protein. Although the numbers were too small for statistical analysis, we noted that mutations 641 T>G, 643 A>G, 647 T>A, and 694 T>C minimally impact 5N inclusion, and all these patients achieved seizure control, while mutations 649 T>C and 660 T>G (both present in Patient #29) increase 5N inclusion, and this patient does not have seizure control.

Notably, Patient #29’s variants T35C/c.649 and C46G/c.660 (using exon/cDNA coordinates) matched 2 of the 3 substitutions that most enhanced exon 5N inclusion (out of the 92 x 3 = 276 substitutions analyzed). Other substitutions in the same positions, as well as substitutions in neighboring nucleotides decreased exon 5N inclusion (Fig. S2). This pattern suggests the presence of ESEs overlapping both positions, with most mutations weakening the ESE motif and the two patient variants uniquely strengthening these ESEs. These results were validated in HEK293 and T98G cells using the mini-gene reporter (Fig. S2f). Analysis of double mutants indicated that patient #29’s combination of T35C/c.649 and C46G/c.660 increased inclusion more than any other combination of mutations in the same or flanking residues, suggesting that this individual may have unusually high expression of mutant protein resulting from very high exon 5N inclusion from the mutant chromosome.

The high frequency of splice-altering mutations in *SCN8A* exon 5N along with the uniformly high third site conservation in *SCN8A* exons 5N and 5A strongly suggests the presence of numerous ESEs or other splicing regulatory elements (SREs) in both exons that could potentially be targeted by ASOs to manipulate splicing. Most known exonic SRE motifs are 4-8 nt long^31,35^, and will thus extend across portions of multiple consecutive codons, with at least half of any individual SRE matching codon positions 1 and 2. Thus, high third site conservation due to presence of conserved SREs is expected to co-occur with additional splicing-related constraint on adjacent first and second positions, which are also highly conserved in these MXEs (Fig. 2a,b, Fig. S3a,b). Consistent with the evolutionary analyses above, we observed a significant correlation between the splicing impact of mutations at different exonic positions and their evolutionary conservation (Fig. 2f, Spearman’s ρ=0.37, *p*=0.0003). Analysis of introns flanking exons 5N and 5A also showed higher evolutionary constraint than in constitutive introns, especially for the intron between exons 5N and 5A (Fig. S3a, S3b). However, even in this intron the fraction of constrained positions was below that observed for third sites in these exons. Together, these observations motivate an ASO screening approach focused on ASOs that directly target exons 5N and 5A rather than the intervening or flanking introns.

**Figure 2:**
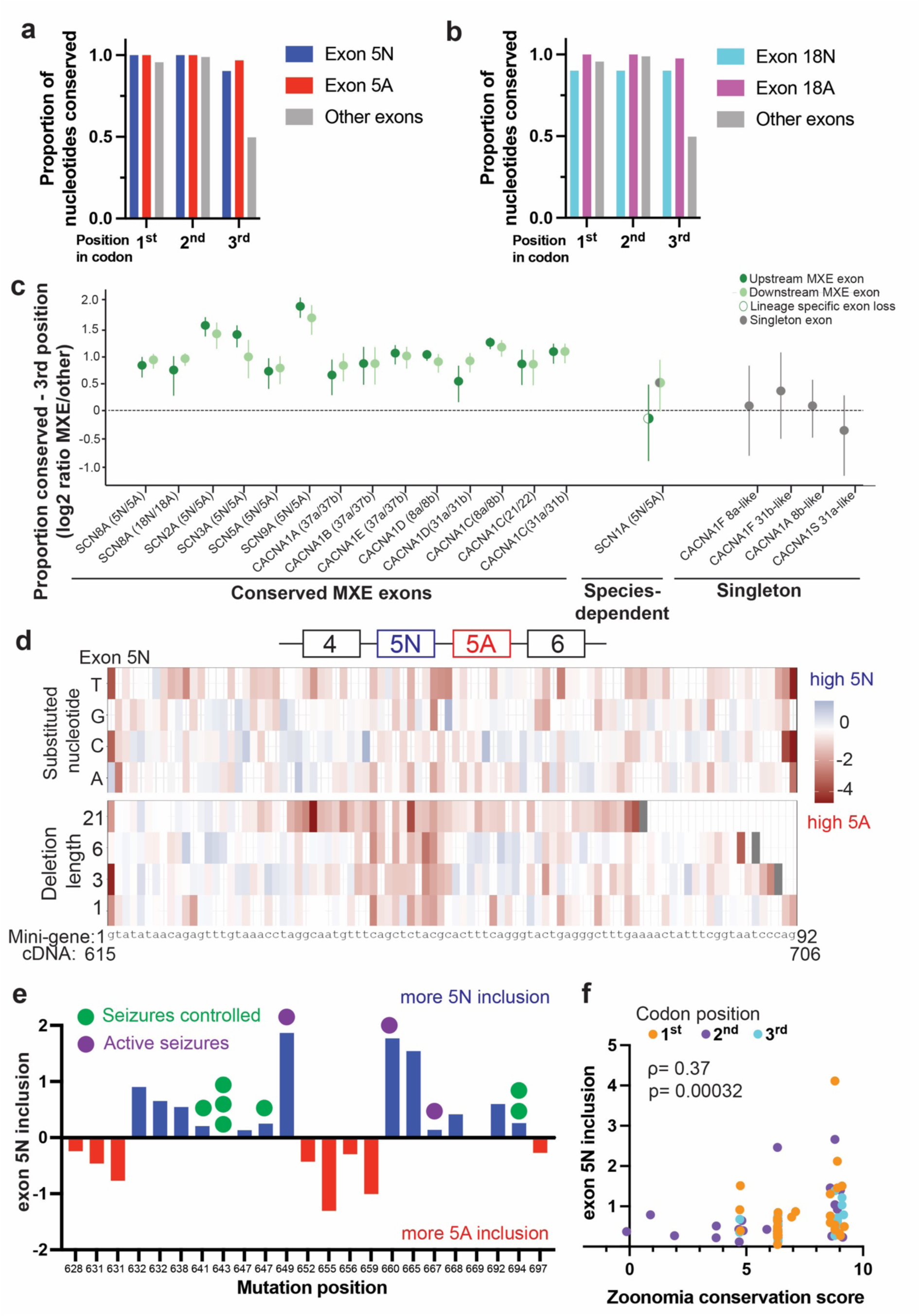
Sequence conservation and splicing regulatory activity of mutually exclusive exons are tightly associated. Plots summarizing the proportion of conserved nucleotides (conservation score ≥2.27) in SCN8A mutually exclusive exons 5N/A **a**) and 18N/18A. **b**). Exon nucleotides are binned by position within codons. **c**) Log2 ratio (MXE/other) of the posterior distributions from our beta binomial model for proportion of codon position three nucleotides, conserved in MXE versus non-MXE in human Na^+^ and Ca^2+^ channel genes. Points represent mean of the log2 ratio of the posteriors (MXE/other). Error bars depict 94% credible intervals. Conserved MXEs are the sets of mutually exclusive exons found in these genes in humans that are conserved across mammals. Species dependent label indicates an MXE set that is mutually exclusive in some mammalian species but not others. Singletons represent the constitutive exons found in *CACNA1F* and *CACNA1S* which have each lost one of the exons in the 8a/b and 31a/b mutually exclusive exon pairs found in the orthologous proteins *CACNA1C* and *CACNA1D*. Note CACNA1C 31a/b is sometimes referred to as 20a/b in other literature. X-axis shows gene (Upstream MXE/Downstream MXE). **d)** Effect on exon 5N enrichment score of individual mutations and deletions of 1, 3, 6 or 21 bp based on SCN8A minigene library. **e)** Known patient mutations and their predicted impact on 5N inclusion based upon the results shown in **d**. **f**) Scatter plot of mean delta fitness score (calculated per position from values in d) versus Zoonomia conservation scores for all positions in *SCN8A* exon 5N. Colors indicate position within a codon for each nt (ρ=0.37, *p*=0003).

### ASOs can modulate the splicing of *SCN8A* exon 5

We sought to identify ASOs that could induce a switch in splicing from 5N to 5A or from 5A to 5N in the mouse *Scn8a* gene which could be used as a therapeutic for patients with exon 5 *SCN8A* mutations by correcting the Na_v_1.6 protein. First, we screened for ASOs that induce a switch from exon 5N to 5A, by tiling exon 5N and adjacent positions (Fig. 3a,b). To measure the 5N:5A ratio in cells, we extracted mRNA, performed RT-PCR, and then digested each sample with two restriction enzymes, each only cleaving one exon (StyI for exon 5N and AvaII for exon 5A). Samples were run on an agarose gel, and the ratio of digested to undigested cDNA was quantified from the image (Fig. S4a). We validated the accuracy of this assay using synthetic cDNAs mixed together at defined ratios (Fig. S4b). For 5N to 5A ASO screening we used mouse neuroblastoma ND7/23 cells, which have high 5N inclusion and lipofectamine transfection (Fig. S4c), using a previously described non-targeting ASO as a control (ASOCTL)^36^. We identified multiple ASOs that modulate *Scn8a* exon 5 splicing, with 11 of 18 ASOs reducing 5N expression, with no changes in *Scn8a* expression (Fig. 3b,c, Fig. S4e). Similar results were obtained in human SH-SY5Y neuroblastoma cells (Fig. S4d). *Scn8a* has high sequence similarity to paralogs *Scn1a*, *Scn2a* and *Scn3a*^37^. ASOs 10-13 had no significant effect on 5N inclusion in *Scn2a* and *Scn3a*, showing selectivity for *Scn8a* (Fig. 3d; note that mouse *Scn1a* lacks exon 5N), while some other ASOs had (undesired) effects on splicing of one or more paralogs and were excluded from further consideration. Primary cortical neurons isolated from the brains of P1 mice and cultured for 7 days had 60% 5N inclusion at baseline, and ASOs 10, 11 and 13, treated via free uptake, reduced 5N levels to less than 10% (Fig. 3e). Effective ASOs were found to align directly to the highly regulatory region of exon 5N identified by mutagenesis, further supporting the presence of multiple ESE elements in this region (Fig. S4g). These data serve to validate 3 ASOs that induce a near complete switch from exon 5N to exon 5A in cultured cells and primary neurons with minimal effects on the splicing of other Na^+^channels or *Scn8a* mRNA expression.

**Figure 3:**
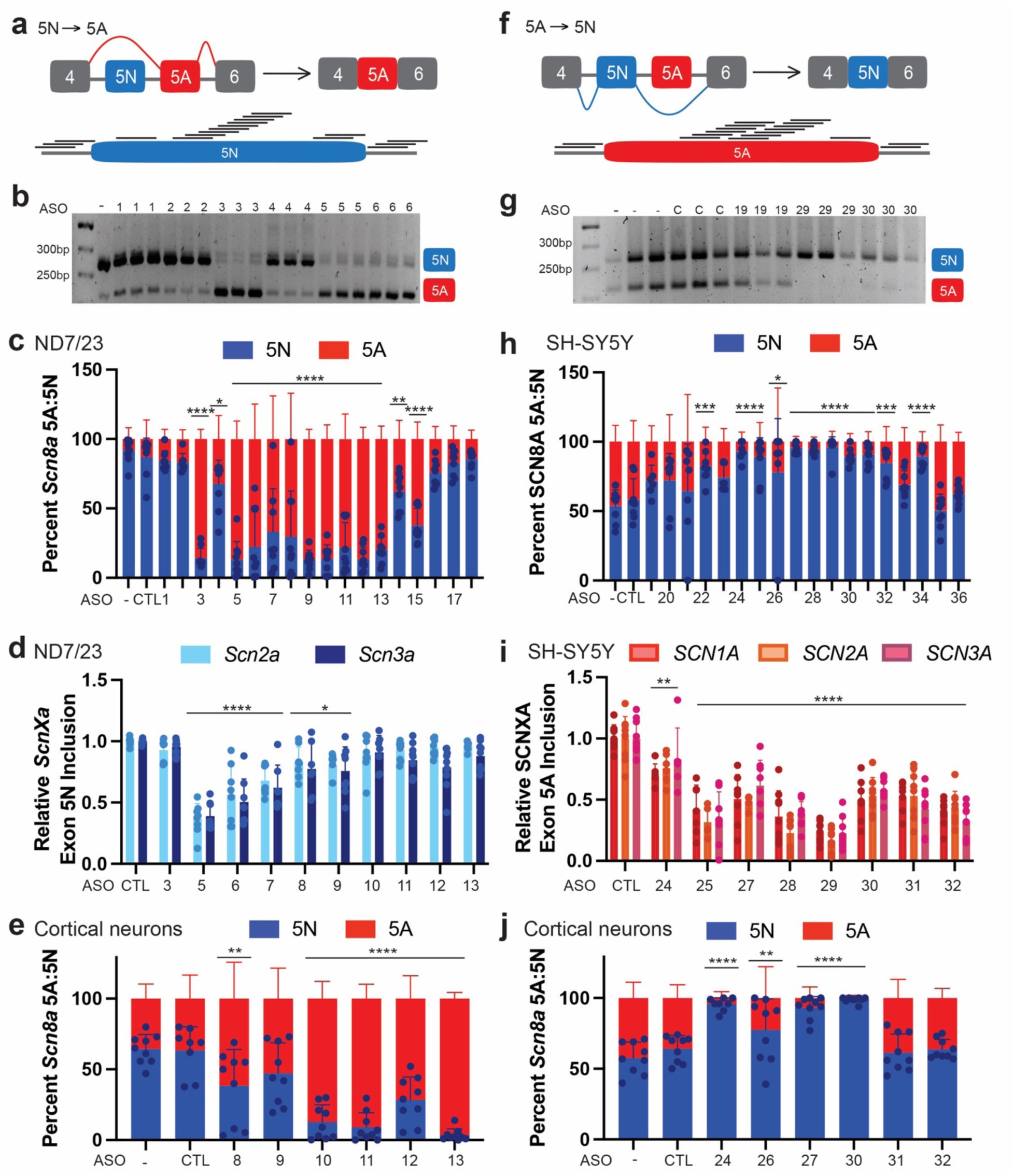
Splice-switching ASOs can be used to regulate the splicing of exon 5 of SCN8A. (**a**) Schematic of ASO intervention to modulate exon 5 alternative splicing to inhibit expression of mutant exon 5N-containing isoform. (**b**) Representative TBE-PAGE of ND7/23 cells treated with 500nM 5N-targeting ASOs. The upper band (280bp) corresponds to the 5N isoform, and the lower band (225bp) corresponds to the 5A isoform. (**c**) Stacked bar plot showing percent SCN8A 5A and 5N inclusion in ND7/23 cells measured after 24hr treatment with 500nM ASO for 24 h by RT-PCR and restriction digestion (**d**) Relative exon 5A:5N inclusion of *Scn2a* and *Scn3a* in ND7/23 cells (**e**) Percent *Scn8a* 5A:5N expression of primary neurons isolated from P0 C57BL/6 pups treated for 24h with 1000nM ASOs. (**f**) Schematic of ASO intervention to modulate exon 5 alternative splicing to inhibit expression of mutant 5A-containing isoform. (**g**) Representative TBE-PAGE of SH-SY5Y cells transfected for 24h with 500nM 5A-targeting ASOs, as in (b). (**h**) Percent SCN8A 5A:5N inclusion in SH-SY5Y measured after 24hr treatment with 500nM ASO for 24 h by RT-PCR and restriction digestion. (**i**) Relative 5A:5N inclusion of *SCN1A*, *SCN2A* and *SCN3A* in in SH-SY5Y cells. (**j**) Percent 5A:5N expression of 1000nM gymnotic uptake ASOs in primary neurons isolated from P0 C57BL/6 pups cultured for 7 days. Data shown as mean ±SD with at least n=4 biological replicates. Statistics by one-way ANOVA, with Dunnett’s multiple comparison test to untreated control, ns P>0.05, * P≤0.05, ** P ≤0.01, *** P≤0.001, ****P≤0.0001.

We next performed a screen for ASOs that could induce a switch in the opposite direction, from 5A to 5N (Fig. 3f). Given the very low baseline inclusion of 5A in ND27/23 cells, we instead used human SH-SY5Y cells, which have just over 50% inclusion of *SCN8A* exon 5A at baseline, taking advantage of the perfect conservation of exons 5A and 5N between human and mouse (Fig. S4c). We identified multiple ASOs that significantly reduce exon 5A inclusion. ASO24, ASO27 and ASO30 reduced exon 5A inclusion in SH-SY5Y cells without impacting *SCN8A* expression, but all had some off-target effects on *SCN1A*, *SCN2A*, and *SCN3A* (Fig. 3g-i, S4f) with ASO24 having the most modest effects. In primary cortical neurons isolated from the brains of P1 mice, 40% inclusion of 5A was observed at baseline, with ASO24, ASO27 and ASO30 reducing 5A levels to less than 10% following ASO treatment (Fig. 3j). These data serve to validate 3 ASOs that induce a near complete switch from exon 5A to exon 5N in cultured cells and primary neurons, with ASO24 having the least off-target effects, and no effects on *SCN8A* mRNA expression. Thus, we identified lead ASOs that can regulate the splicing of exon 5 of *SCN8A* in either direction.

### Splice-switching ASOs rescues phenotypes in patient-derived iPSC-derived neurons

We next explored whether these ASOs have therapeutic potential for patients with variants in *SCN8A* exon 5. Specifically, we focused on patient #29, who has 2 de novo variants in exon 5N, located in *cis*: 649 T>C, S217P, which changes a residue in the extracellular loop, and the synonymous variant 660 C>G, R220R, which does not impact Na_v_1.6 activity. These two variants substantially increase exon 5N inclusion (Fig. 2, Fig. S2). This patient has a severe phenotype, with uncontrolled seizures presenting as infantile spasms, tonic, clonic and tonic-clonic seizures, severe global developmental delay, and hypotonia^19,38^. We isolated fibroblasts from this patient and generated induced pluripotent stem cells (iPSCs), confirming these variants by sequencing and karyotyping (Fig. S5a,b). We differentiated these cells into cortical-like (NGN2) iPSC-neurons using an established tetracycline(tet)-inducible system^39,40^, and confirmed expression of Na_v_1.6 (Fig. S5c,d). Using a multielectrode array (MEA) to extracellularly measure spontaneous electrical activity^40,41^, we found that mature S217P^+/-^ iPSC-neurons show increased neural network activity compared to a sex-matched wildtype control embryonic cell line, with significant increases in mean firing rate, bursting frequency and burst duration (Fig. 4a-e). We then evaluated the splicing dynamics of *SCN8A* exon 5 at major differentiation milestones: induction, maturation and maintenance^39,40,42^ (Fig. 4f,g). Exon 5N inclusion decreased from ∼90% to ∼10% during differentiation of control iPSCs. By contrast, in S217P^+/-^ iPSC-neurons 5N remained elevated, persisting at 50-60% levels in mature neurons after DIV 14 (Fig. 4f,g). Because transcripts from the reference allele in S217P^+/-^ iPSC-derived neurons are presumably regulated similarly to those in control iPSC-derived neurons, the elevation of 5N levels is likely driven exclusively by inclusion of transcripts containing the two de-novo variants.

**Figure 4:**
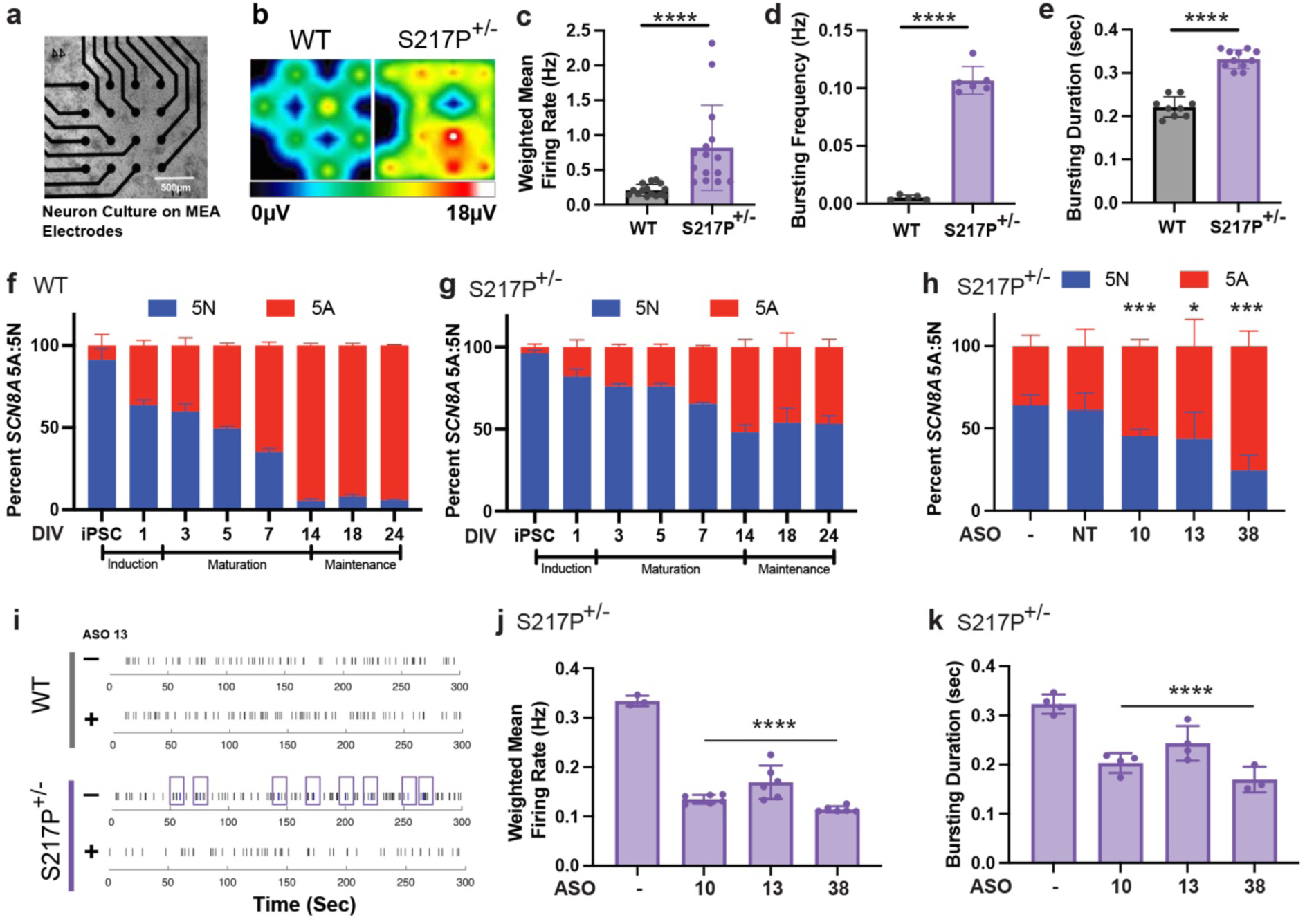
Mutant S217P^+/-^ iPSC-derived induced neurons recapitulate splicing switch delay, and phenotype can be rescued by ASO treatment. **a**) MEA well plate with iPSC-derived S217P^+/-^ cultures at DIV14 before recording. **b**) Representative heat map of neuronal activity on 4x4 grid of MEA at DIV18. Field-potential (µV) color-coded as shown in color bar below. (**c**) Weighted mean firing rate, (**d**) bursting frequency, and (**e**) bursting duration based on quantification of MEA neuronal network parameters. Data were pooled from biological replicates (wells) across three separate differentiations, with *n* = 15 for control and *n* = 15 for S217P^+/-^ cultures. **f,g**) Relative percent endogenous SCN8A 5A:5N expression through differentiation of inducible iPSC-neurons for control and S217P^+/-^ neurons. **h**) Relative percent SCN8A 5A:5N expression in DIV18 S217P^+/-^ iPSC-neurons 5 days after 500nM ASO treatment. **i-k**) (**i**) representative bursting events (purple boxes) in raw spiking activity, (**j**) weighted mean firing rate in S217P^+/-^ cultures, and (**k**) bursting duration in S217P^+/-^ cultures at DIV18 5 days post ASO treatment (500nM transfection). Data for S217P^+/-^ ASO treatments were pooled from biological replicates (wells) across two separate differentiations; the sample size was *n* = 7 (wells) for untreated control and *n* = 6 (wells) for ASO treatments. For all data, each point is the mean of at least three technical replicates per biological sample, with error bars representing ±SEM. Student’s *t*-test was performed when data were normally distributed (by Kolmogorov-Smirnov normality test), and Mann–Whitney U was performed when data were not normally distributed all relative to untreated control; ns p>0.05, *p ≤ 0.05, **p ≤ 0.01, ***p ≤ 0.001, ****p≤0.0001.

We next determined the efficacy of our top ASO candidates in the mutant iPSC-derived neurons. We treated mature iPSC-neurons for five days with 500nM ASO10, ASO13, or a similar ASO with sequence complementary to the patient mutation, ASO38, and found that all 3 ASOs reduced relative exon 5N expression in iPSC-neurons compared to non-targeting and untreated controls (Fig. 4h, Fig. S5e). Further ASO treatment functionally rescued S217P^+/-^ iPSC-neuron physiological activity. Temporal visualization of bursting activity by raster plots indicated a reduction in S217P^+/-^ bursting numbers to near control levels (Fig. 4i). All three targeting ASOs restored weighted mean firing rate and bursting duration in S217P^+/-^ iPSC-neurons to control levels (Fig. 4j,k). Wildtype iPSC-neuron activity remained unaffected by ASO treatment (Fig. S5f,g). These data demonstrate the potential of these ASOs to restore normal activity in neurons of patients with *SCN8A* exon 5N mutations.

### Novel mouse model of SCN8A DEE with patient exon 5N variants has seizures and motor impairment

We generated a novel mouse model of SCN8A DEE containing patient #29’s exon 5N S217P and R220R variants, designated S217P^+/-^. We characterized the first generation of mice (F1) generated from mutant embryos, which lived until adulthood (Fig. S6a). F1 S217P^+/-^ mice presented with spontaneous seizures, confirming an epileptic phenotype as expected. All adult (10-13 week-old) mice that were implanted with bilateral ECoG electrodes and continuously recorded for 14 days experienced seizures. The average seizure burden per mouse was 1.2 seizures/day with a mean seizure duration of 42.1 seconds ± 18.7 seconds (Fig. S6b-c). Behavioral analysis revealed that F1 S217P^+/-^ adult mice spend less time actively moving in an open field and additionally have reduced vertical ambulation, suggesting a motor impairment (Fig. S6d-e). F1 animals have significantly higher inclusion of 5N, up to 45%, from 7-13 weeks of age compared to WT mice of similar age which have only 5% inclusion, though at 36 weeks mutant levels are similar to WT (Fig. S6f). We did not see any changes in neuronal number as quantified by NeuN staining in the cortex. However, adult F1 S217P^+/-^ mice have reduced numbers of parvalbumin-positive interneurons, a phenotype previously described in other models of genetic epilepsy but not for SCN8A variants^43,44^ (Fig. S6g-h). We concluded that our F1 generation constitutes a model of genetic epilepsy with the animals not only having seizures, but additionally changes in motor behavior, splicing and interneuron populations. The high exon 5N inclusion in this model, consistent with the mutagenesis and iPSC data above (Fig. 2d, 4g), makes this model more challenging to rescue with ASOs, suggesting that it provides a particularly stringent assessment of ASO efficacy.

The F2 S217P^+/-^ generation of mice displayed a severe DEE-like phenotype and profoundly reduced survival, with a median survival of just 21 days (Fig. 5a). Onset of pathology is seen at P15 with a significant reduction in weight compared to littermates that continues until death (Fig. 5b). Up until P15 there is no discernable phenotype in the mutant mice, based on neonatal behavioral assays and righting reflex (Fig. 5c). Seizure-like behavior from daily video recordings was quantified using a previously published adjusted Racine Scale (1-6) for pups^45–47^, with seizures starting at P15. F2 pups on average experience >5 seizures during 10 minutes of quantification (Fig. 5d). At P16 the most severe seizures seen were scored 4-6 on the adjusted Racine Scale, with P15-P17 seizures lasting an average of 24 seconds (Fig. 5e-f). Along with the seizure phenotypes, the pups also exhibit changes in ambulatory ability. S217P^+/-^ pups experience numerous falls at P15, which increase over time (Fig. 5g). At the same time, mutant pups start spending extended time on their back or side. From P18-19 pups were unable to move or hold themselves up, at which point they were sacrificed (Fig. 5h). P0 and P13 S217P^+/-^ mice have significantly higher inclusion of *Scn8a* 5N (50%) compared to wild-type littermates (10%), confirming that this mutation increases exon 5N splicing (Fig. 5i). It is not clear why the F2 generation is more severely impacted than the genetically identical F1 generation, whether because of parental effects, epigenetics, or environmental differences. Overall, the S217P^+/-^ mutant mice demonstrate the rapid onset of disease progression at P15 once seizures start. Notably, reduced survival in this model does not seem to result from seizures, with death not observed proximal to seizure occurrence, but appears to occur through the rapid deterioration of movement ability post-seizure onset, impacting the pups’ ability to feed, leading to weight loss and starvation. This phenotype is reminiscent of the clinical presentation of SCN8A DEE patients, with 90% of patients experiencing hypotonia which can lead to a number of comorbidities which contribute to disease burden and early mortality.

**Figure 5:**
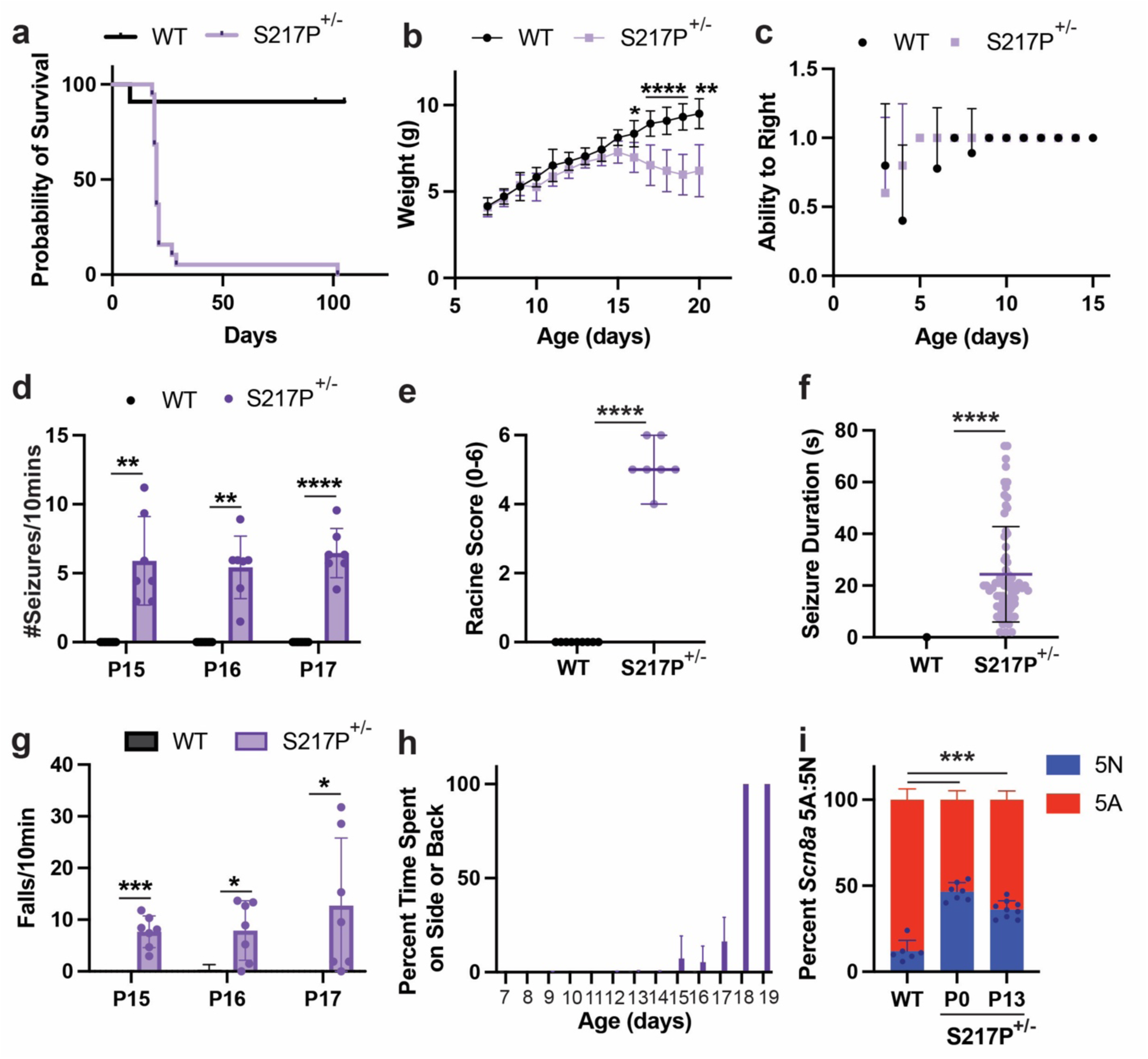
F2 S217P^+/-^ have a severe DEE phenotype with reduced survival, seizures and motor phenotype. **a**) Survival of WT and S217P^+/-^ animals. Survival was monitored for 110 days, *n* = 30 (WT = 11, S217P^+/-^ = 19). **b**) Weight was tracked daily for treated animals from P7-P20, *n* = 37 (WT = 18, S217P^+/-^ = 19). **c**) S217P^+/-^ mutants and WT littermates’ ability to right was observed from P3-P15, score as 1 = able to right, 0 = unable to right, n = 37 (WT = 18, S217P^+/-^ = 19). **d-g**) Quantification of P16-17 video analysis for (**d**) seizure frequency, n = 16 (WT = 9, S217P^+/-^ = 7), (**e**) P16 Racine scoring, *n* = 16 (WT = 9, S217P^+/-^ = 7), (**f**) P15-P17 seizure duration *n* = 72 (WT = 1, S217P^+/-^ = 71) and (**g**) P15-P17 falls, n = 16 (WT = 9, S217P^+/-^ = 7). **h**) Percent time S217P^+/-^ and WT (P14-P20) mice spend on their back or side from video analysis, *n* = 16 (WT = 9, S217P^+/-^ = 7). (**i**) *Scn8a* 5A:5N quantification of S217P^+/-^ (P0 and P13) and WT (P0) whole brains, *n* = 22 (WT P0 = 6, S217P^+/-^ P0 = 7, S217P^+/-^ P13 = 9). Statistics: (**b**) multiple unpaired t tests, (**d-f**) One sample t and Wilcoxon test, theoretical mean set to 0, (**i**) one-way ANOVA with Dunnett’s multiple comparison test to WT P0, ns P>0.05, * P≤0.05, ** P ≤0.01, *** P≤0.001, ****P≤0.0001.

### Treatment with a 5N splice-switching ASO improves seizures, motor impairment and survival of S217P^+/-^ mutant mice

To assess safety of ASOs *in vivo*, 8-12 week-old WT mice were treated with a single high 700 µg dose of ASO-CTL, ASO10, ASO11 or ASO13 via intracerebroventricular (ICV) injection and monitored for adverse events following the treatment. At 3 and 24h, ASO-CTL and ASO13 had no adverse events associated with toxicity, such as hunching, stiffness, hyperactivity or seizures^48,49^. ASO10 and ASO11 had mild adverse events. Additionally, there was no difference in weight compared to PBS-treated controls over 8 weeks (Fig. S7a-c). To assess changes in immune response post treatment, the expression of GFAP and CD68 were measured, with mean GFAP and CD68 mRNA levels not increased for any ASO (Fig. S7d-e). Based on these safety data, and efficacy and off-target data presented above, we prioritized ASO13 for subsequent experiments. To assess long-term behavioral impacts of ASO treatment in wild-type C57BL/6 mice, pups were treated with 60 µg ASO via ICV injection at P1. Pups’ weight, ability to right and performance on negative geotaxis was assessed daily from P7 to P14, with no significant differences seen between untreated, ASO-CTL and ASO13-treated animals (Fig. S8a,b). Starting at 6 weeks, adult mice underwent behavioral tests (Open-Field, Y-maze and Light/Dark box) to determine if there were impacts on motor ability, anxiety-like behavior or spatial working memory. No significant differences were observed between untreated adult mice and mice treated with ASO-CTL or ASO13 (Fig. S8c-p). Together these results indicate a favorable safety profile for ASO13.

To assess the therapeutic potential of ASO13, we treated F2 S217P^+/-^ mice with 60 µg of ASOCTL or ASO13 via ICV injection at P1. The pups were monitored daily for weight (P4+), righting reflex (P7-P14), negative geotaxis (P7-P14) and video recording (P15-P20), and surviving adult mice underwent Open Field and Y-maze tests (6w), with sacrifice at 10-12w followed by RNA splicing analysis and immunohistochemistry (Fig. 6a). ASO13 significantly improved survival of S217P^+/-^ mice, from 0% survival to P30 for pups treated with ASOCTL to 30% survival to P30 with ASO13 (Fig. 6b). In S217P^+/-^, ASO13 treatment reduced the inclusion of exon 5N from 45% to 10% (Fig. 6C). ASO13 had no effect on pup behavior in either WT or mutant mice (Fig. S9a-b). ASO13 substantially reduced the number and the severity of seizures in S217P^+/-^ mice (Fig. 6d,e). Further, it significantly reduced the number of falls (Fig. 6f), indicating that this treatment not only benefits seizures, but also improves motor function.

We next compared S217P^+/-^ mice that died by P21 versus those that survived. Both ASOCTL and ASO13-treated S217P^+/-^ mice lost weight around weaning age (P16-21), though surviving ASO13-treated animals returned to WT weight levels by 6 weeks of age (Fig. 6g). No significant differences between levels of 5N inclusion, seizure prevalence, severity or number of falls were observed in the survivor population versus ASO13-treated S217P^+/-^ mice that died (Fig. S9 c-f). However, ASO13-treated S217P^+/-^ animals had reduced time spent on their back or side compared to ASOCTL-treated S217P^+/-^ pups, with ASO13-treated surviving mice not spending any time on their back or side, similar to WT mice (Fig. 6h). These observations suggest that improvement in motor function/hypotonia, not just seizures, contributes to survival to adulthood of ASO13-treated mice. ASOCTL and ASO13-treated mice that died (average age of death = P19), had fewer parvalbumin-positive interneurons in the cortex compared to WT treated animals. Interestingly, ASO13 survivors had equivalent numbers of parvalbumin-positive interneurons at 10-13 weeks of age compared to age-matched WT ASO13-treated animals (Fig 6i-k). There were differences in NeuN labeled cell density in the cortex across groups at P19, with a slight decrease in NeuN-positive cells in the mutant ASO13-treated survivors relative to WT age-matched ASO13-treated controls (Fig. S9g-i). At P19, ASO13 intensity was measured throughout the brain with robust distribution of the ASO in the cortex, hippocampus, thalamus and hypothalamus in WT and S217P^+/-^ animals (Fig. S9j-k). Thus, successful ASO13 treatment in S217P^+/-^ improves survival, motor function, seizure occurrence and prevents loss of cortical interneurons.

**Figure 6:**
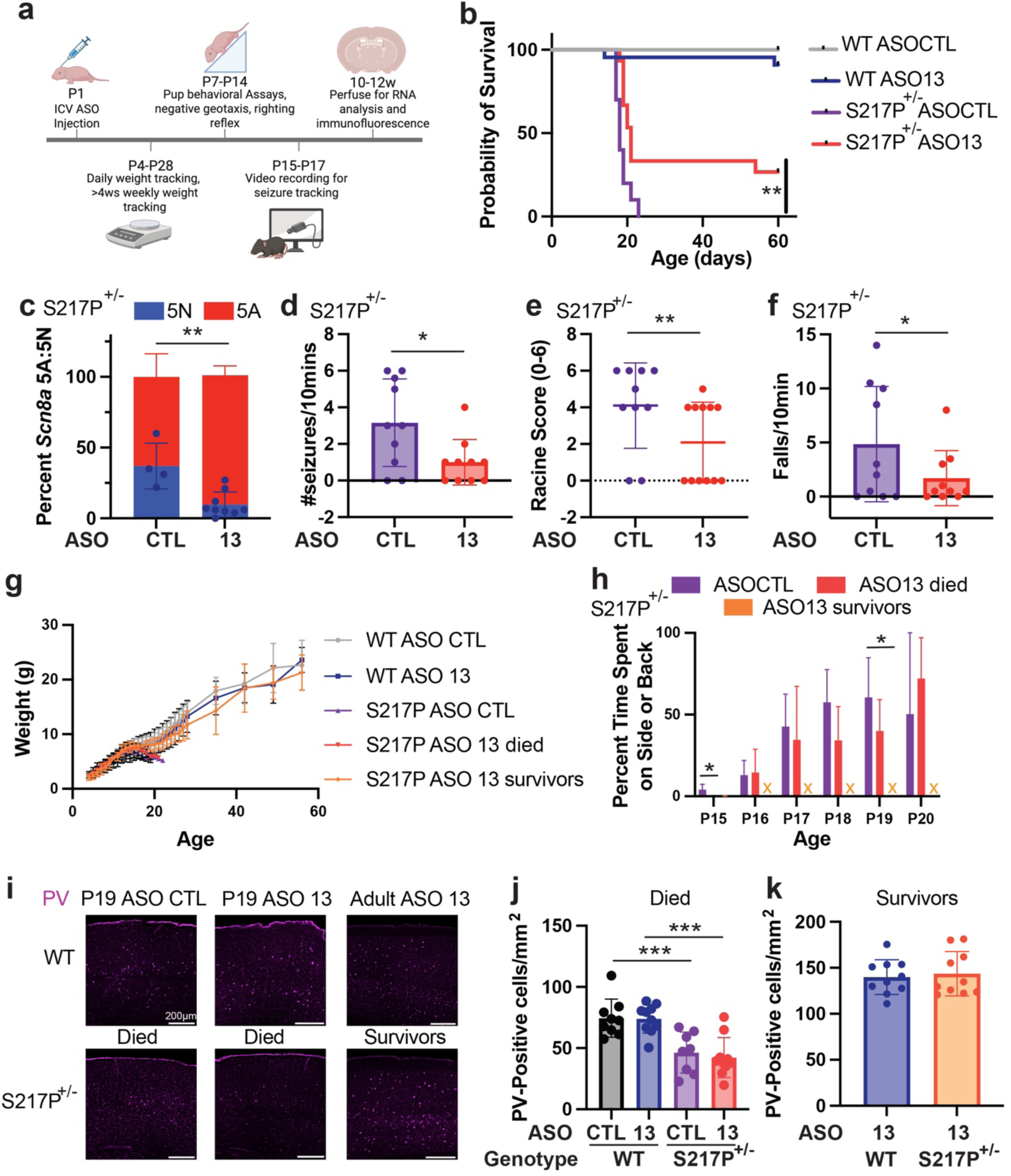
ASO-13 treatment improves survival, seizure burden and motor impairment in S217P^+/-^ mice. **a**) Experimental plan for ASO treatment of S217P^+/-^ mice. Mice received P1 ICV injections of 60µg ASO. Weight tracking (P4-P60), behavioral assessment (P7-P14) and video recording (P15-P20) was conducted throughout development. RNA was isolated at time of death (P17-23) or at end point (10-13w) with immunofluorescence. **b**) Survival of ASO-treated WT and S217P^+/-^ animals. Survival was monitored for 60 days, *n* = 37 (WT ASO-CTL = 10, WT ASO-13 = 22, S217P^+/-^ ASO-CTL = 10, S217P^+/-^ ASO-13 = 15). **c**) *Scn8a* 5A:5N quantification of S217P^+/-^ ASO-CTL and ASO-13 treated animals at death (P16-P23) or endpoint (10-13w), *n* = 13 (CTL = 4, ASO-13 = 9). **d-f**) Quantification of P16 video analysis for seizure frequency (**d**), Racine scoring (**e**) and falls (**f**), *n* = 10 animals per group. **g**) Weight was tracked daily for treated animals from P4-P28, then weekly from P28-P60, *n* = 38 (WT ASO-CTL = 10, WT ASO-13 = 23, S217P^+/-^ ASO-CTL = 10, S217P^+/-^ ASO-13 died = 10, S217P^+/-^ ASO-13 survived = 5). **h**) Percent time ASO treated S217P^+/-^ (P15-P20) mice spend on their back or side from video analysis, *n* = 20 (S217P^+/-^ ASO-CTL = 10, S217P^+/-^ ASO-13 died = 7, S217P^+/-^ ASO-13 survived = 3). (**i**) Representative images of Parvalbumin (PV) positive cells, pink, across groups. Scale bar, 200um. (**j**) Quantification of PV positive cells/mm^2^ in ASO treated WT and S217P^+/-^ animals that were sacrificed at mean on P19, *n* = 18 animals (WT ASO-CTL = 4, WT ASO-13 = 5, S217P^+/-^ ASO-CTL = 5, S217P^+/-^ ASO-13 = 4, 1-3 slices/animal). (**k**) Quantification of PV positive cells/mm^2^ in ASO treated WT and S217P^+/-^ adult surviving mice (10-13 weeks old), *n* = 7 animals (WT ASO-13 = 4, S217P^+/-^ ASO-13 survivors = 3, 1-4 slices/animal). Statistics – (**b**) Mantel – Cox test, (**c-f**) unpaired *t*-test, (h) Kruskal-Wallis test with Dunn’s multiple comparison to S217P^+/-^ ASOCTL. (**j-k**) Linear mixed modeling, comparing genotype and treatment, slices are technical replicates to biological animal, ns P>0.05, * P≤0.05, ** P ≤0.01, *** P≤0.001, ****P≤0.0001.

## Discussion

There is an unmet need for novel therapeutics for children suffering from DEEs, which not only cause intractable epilepsy but also a multitude of non-seizure comorbidities, often driven by hypotonia, which have devastating impacts on quality of life and can contribute to early death. Here, we sought to improve treatment of SCN8A epilepsy/DEE by developing a disease-modifying ASO treatment that shifts cellular production away from pathogenically altered neonatal isoform (5N) mRNA toward production of wildtype mRNA of the adult (5A) isoform, without changing overall *SCN8A* expression. We identified the alternatively spliced exon 5 of *SCN8A* as a critical region of the gene, where pathogenic variants lead to severe disease, characterized by a high incidence of infantile spasms, and where patient outcome may be impacted by gradual shift in splicing from exon 5N to 5A. We have developed splice-switching ASOs that substantially shift splicing from exon 5N to exon 5A, or vice versa. Using iPSC-derived neurons from a patient with an exon 5N S217P^+/-^ mutation, we show that splice-switching ASOs reduce exon 5N inclusion and spontaneous spiking. In a novel mouse model of SCN8A exon 5N S217P^+/-^, splice-switching ASOs can reduce seizures, improve motor function, and substantially extend lifespan.

Current therapies for patients with SCN8A and other severe pediatric epilepsies primarily use small molecule drugs to reduce excitatory and/or increase inhibitory synaptic activity, with seizures being the most common primary endpoint in clinical trials. No approved treatments are demonstrated to improve the non-seizure comorbidities that are so prevalent in patients with severe DEEs and reduced lifespan. Here, we introduce a model of pediatric epilepsy which demonstrates both seizures and impaired motor function, with early death attributable to the inability to move and therefore eat, similar to what is seen in patients who often require feeding tubes for nutrition. Our splice-switching ASOs reduced the production of pathogenically altered mRNA without affecting total *Scn8a* mRNA in the CNS, representing a more precise and potentially effective therapy that both reduced seizures and improved motor development and other comorbidities. The mechanisms by which an overactive CNS leads to hypotonia, motor impairments and impaired peripheral organ function are poorly understood. However, our data demonstrates that correcting the source of the disorder in the CNS can improve both seizure and non-seizure outcomes, which highlights the potential of genetic therapies for DEEs caused by pathogenic variants in ion channels.

Our data also highlight the importance of understanding the degree of gene correction required to obtain meaningful clinical impacts. Using a conditional allele of an *Scn8a* R1872W mutation, the Meisler lab found that expression of just 8% of mutant *Scn8a* transcript was sufficient to cause spontaneous seizures^50^. This suggests that high levels of genetic correction are needed to obtain seizure freedom, at least in mice, which do not receive other clinical interventions that patients receive^50^. Consistent with a requirement for high levels of genetic correction, we observed 30% survival of treated mice, with significant reductions in seizures and complete rescue of the motor phenotype after a single ASO injection. The ASO reduced mutant exon 5N inclusion substantially, from over 50% to ∼10%. Compared to treated mice that died by day 21, treated mice that survived to adulthood showed lower exon 5N inclusion (6% compared to 13%), suggesting that higher correction could improve outcome, though this difference was not statistically significant (Fig. S9). Further optimization of dosing, chemistry and formulation through standard pharmacology are likely to further improve the therapeutic potential of these approaches. Alternate genetic therapies have been tested in two other SCN8A mouse models, both of which have seizures and early mortality but lack the severe motor phenotype observed in our S217P^+/-^ model. In both cases, a roughly 30% reduction in mutant transcripts was sufficient to reduce seizures and improve survival^51,52^. These observations suggest that different levels of genetic correction may be required depending on the severity of the mutation, with high levels of genetic correction required to correct the severe seizure and motor phenotype seen in our model.

Pipelines for clinical development of ASOs rely on the identification of a large number of candidates with reasonable efficacy, which are then screened for off-target effects and toxicity while optimizing efficacy. The uniformly high conservation of *SCN8A* exons 5N and 5A across all three codon positions (Fig. S3a,b) and the large fraction of substitutions that altered minigene splicing imply a high density of SREs in these exons, likely explaining the high yield of splice-switching ASOs from our screening relative to other ASO tiling studies^53–55^. The observation of similarly elevated third position conservation across Na^+^ and Ca^2+^ channel MXEs suggests that their splicing may be similarly responsive to exon-targeting ASOs, with the potential for identification of large numbers of efficacious lead ASOs. Patients who have variants in exon 5 of other CNS-expressed sodium channel genes, SCN1A and SCN2A, have been reported, creating the opportunity for development of novel ASO therapeutics using an approach similar to that described here^56–59^. While direct targeting of exons appears most promising, the high similarity of these exons to exons in paralogous VGSCs (Fig. S10a,b) suggests that some tested ASOs may exert “off-target” effects on the splicing of other sodium channels, as seen for a subset of ASOs tested here (Fig. 3d,3i). The intron between exons 5N and 5A is quite conserved between orthologs, but has much lower sequence similarity between paralogs, providing an alternate target region for ASO screening with reduced off-target potential. Splice switching of coding MXE exons is expected to minimally impact gene expression^53^, which may provide a potential therapeutic advantage over most other ASO approaches for applications to dosage-sensitive “Goldilocks” genes such as the genes studied here, and help bring ASO treatments to a broader population.

## Materials and Methods

### Patient data

52 patients were identified from the literature^1–13^ and International SCN8A registry^14^. Patient information is in Table S1. The seizure onset, seizure incidence and infantile spasm incidence data for the exon 5 population was compared to the entire published SCN8A patient population ^15,16^. The location of each mutation was mapped to each exon based on chromosome coordinate data: Exon 5N corresponds to positions 51,688,758 to 51,688,849 for hg38 and 52,082,542 to 52,082,633 for hg19. Exon 5A corresponds to positions 51,689,005 to 51,689,096 for hg38 and 52,082,789 to 52,082,880 for hg19. The incidence of infantile spasms was calculated from the 270 individuals published in Figure S1 of ^16^, where 64 were reported to have experienced infantile spasms. Of 12 individuals with variants mapping within exon 5N/5A, 9 were reported to have experienced infantile spasms, while 3 did not. We performed a binomial test to determine whether an excess of IS patients had variants in exon 5 compared with the 55 IS patients with variants mapping in the remainder of the Na_V_1.6 channel).

### Saturation Mutagenesis and Deletion Analysis

#### Construction of pCMV3 SCN8A 4-5N-5A-6 minigenes

A minigene was designed to study alternative splicing of exons 5N and 5A of the human SCN8A gene. The minigene contained the complete sequences of exons 4, 5N, 5A and 6, the complete sequence of intron 5N, and ∼200 base pairs of the 5’ and 3’ ends of introns 4 and 5A, cloned in expression vector pCMVBeta downstream of a CMV promoter and upstream of a SV40 polyadenylation signal. Truncation of introns 4 and 5A was necessary because the length of these introns precluded cloning of the entire genomic sequence in the expression vector. The different segments of SCN8A gene were PCR amplified from human genomic DNA (PROMEGA, G3041) and assembled by Gibson cloning. The following primers were used:

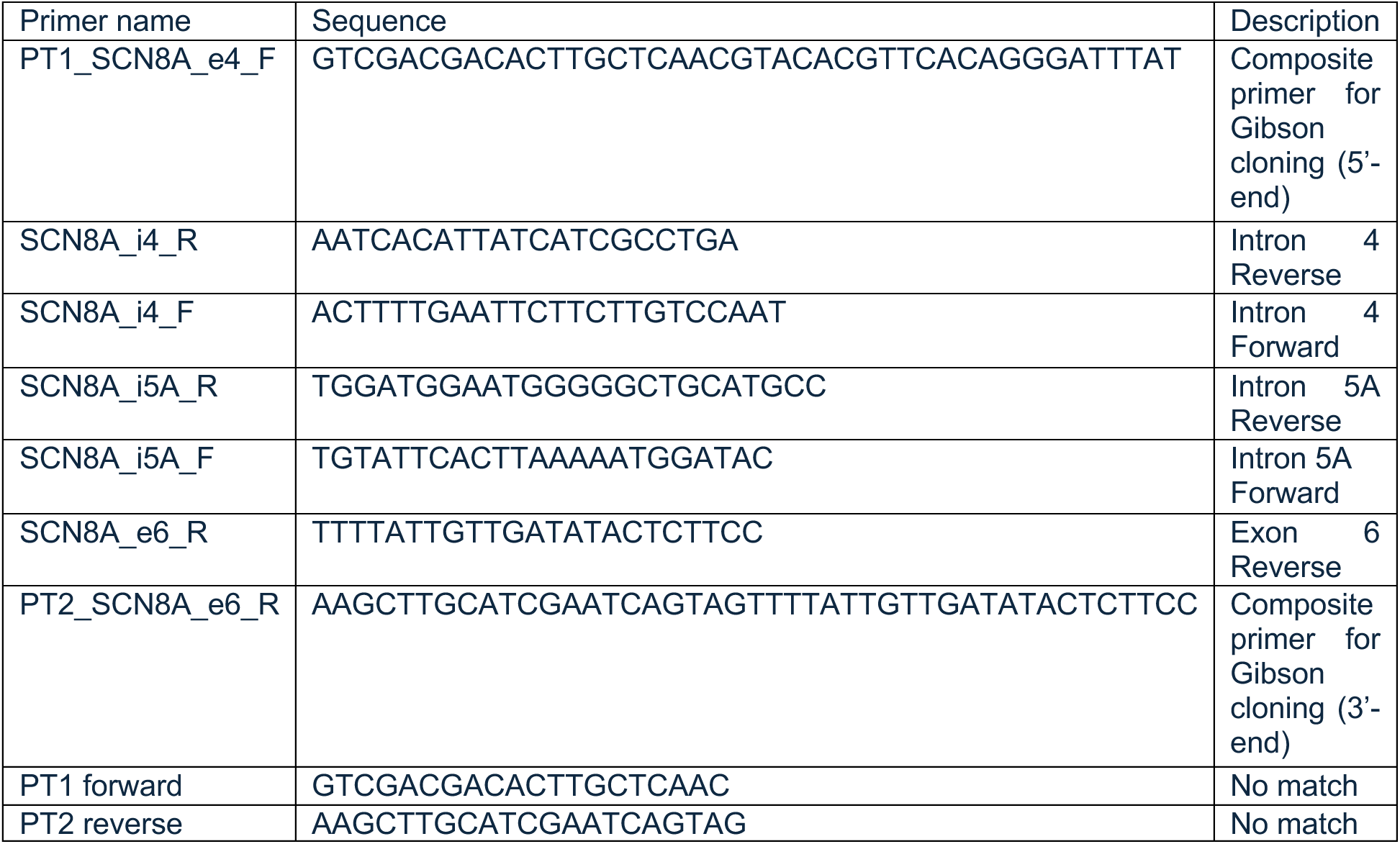

The primers corresponding to the 5’ end of exon 4, and 3’ end of exon 6, contained 5’ extensions (PT1, PT2) corresponding to two sequences without any match to the human genome or transcriptome, to facilitate the amplification of transcripts derived from the minigene construct ^17^ and not from endogenous SCN8A transcripts. The wild type minigene was subsequently modified by PCR-mediated mutagenesis to generate the following minigene variants, which were verified by Sanger sequencing:

| Minigene | Sequence in exon 5N |
| --- | --- |
| pCMV SCN8A e4-e5N-e5A-e6 | Wild type |
| pCMV SCN8A e4-e5N-e5A-e6 T35C | T35C (c.649T->C / p.S217P) |
| pCMV SCN8A e4-e5N-e5A-e6 C46G | C46G (c.660 C>G / p.R220R) |
| pCMV SCN8A e4-e5N-e5A-e6 T35C/C46G | Both T35C and C46G |

Additional mutants were generated to validate the results of the saturation mutagenesis library (see below). Two independent sequence-verified clones for each mutant design were tested in transfection assays, with equivalent results.

#### Cell transfections and analysis of spliced isoforms

Minigenes were transfected in three technical triplicates, in human embryonic kidney Hek293, cervical cancer HeLa, glioblastoma multiforme T98G and neuroblastoma SH-SY-5Y cell lines. 30 nanograms of plasmid DNA were transfected onto 200,000 cells at 70% confluency in 6-well plates, plated 12 hours earlier, using Lipofectamine 3000 (Thermo Fisher Scientific, L3000015) for T98G cells, or Lipofectamine 2000 (Thermo Fisher Scientific, 11668019) for Hek293 cells, and OPTI-MEM reduced serum media (Thermo Fisher Scientific, 31985070). RNA was isolated 48 hours post-transfection using Maxwell RSC SimplyRNA Cells (PROMEGA, AS1390). 200 - 400 ng of total RNA were reversed-transcribed with Superscript III reverse transcriptase (Thermo Fisher Scientific, 18080085) using 2 picomole vector-specific PT2 reverse primer in a 20 µliter reaction. PT2 reverse 5’ AAGCTTGCATCGAATCAGTAG-3, PCR amplification was performed using 1.5 µliters of cDNA (from the 20 µl RT reaction) and 15 picomoles of PT1 and PT2 vector-specific primers in a 30 µl reaction with GoTag flexi G2 DNA polymerase system (PROMEGA, M7806).

| PCR program |  |
| --- | --- |
| 95°C | 2 minutes |
| 95°C | 15 seconds |
| 55°C | 30 seconds |
| 72°C | 60 seconds |
| Goto step 2 | 29 cycles |
| 72°C | 5 minutes |
| 10°C | forever |

DNA amplification products were resolved by electrophoresis on non-denaturing 6% acrylamide gels. Representative results are shown in Figure S2 for wild type and a variety of mutant constructs, as indicated. The expected sizes of the amplification products for each isoform (including an isoform that includes both exons 5N and 5A, which is also detected at low proportion (4%) in GTEX) are indicated, along with the relevant molecular masses of the markers (GeneRuler 50 bp, Thermo Fisher Scientific). Minigene transfections and RT-PCR analyses were carried out in at least 3 biological replicates, each with 3 technical replicates, with consistent results.

In HEK293 cells there are three main products of amplification (Fig S2), which correspond in size to the inclusion of the two alternative exons (5N+5A), the inclusion of a single product (5N or 5A, which are of the same length -92 bp-are therefore indistinguishable) and the product of exon skipping. Roughly equal amounts of double and single exon inclusion product are observed in Hek 293 cells, and lower amounts of double skipping products. Comparison of the products of RT-PCR of wild type and mutants in HEK293 cells indicates that the relative abundance of the double inclusion band (5N+5A) increases with mutants T35C, C46G, T35C+C46G (mutations found in the patient), as well as T51C and T78C (mutations tested to validate the results of the high-throughput mutagenesis, see below), while it decreases with mutants G45T, C46T and T78G (another set if mutants tested to validate the results of the high-throughput mutagenesis, see below) in favour of the products of single exon (5N or 5A) inclusion.

The pattern of products in T98G cells is a bit different from that of HEK293 cells, as the product corresponding to double exon inclusion (5N+5A) is by far the most abundant, with much less single exon inclusion product and barely any detectable double skipping product. There is an additional product of amplification slightly above 250 bp which was not detected in HEK293 cells and that sequencing determined to correspond to vector sequences. As observed in HEK 293 cells, however, comparison of the products of RT-PCR of wild type and mutants indicate that the relative abundance of the double inclusion band (5N+5A) also increases (even more) in T98G cells with mutants T35C, C46G, T35C+C46G, T51C and T78C, while it decreases in mutants G45T, C46T and T78G in favour of the products of single exon (5N or 5A) inclusion. Similar results were obtained in HeLa and SH-SY-5Y cell lines.

As mentioned above, as the length of exons 5A and 5N is identical (92 nucleotides), it is not possible to distinguish between the products of inclusion of the two isoforms by the size of the amplification product alone. To assess the relative use of exons 5A and 5N in wild type and mutant minigenes, two strategies were followed.

In the first strategy, products of amplification of single (5A/5N) exon inclusion were gel isolated and cloned in pCMVBeta, and 12 individual clones per variant under study were Sanger-sequenced. While some clones did not provide readable sequences, the following table summarizes the results obtained for the clones that could be assigned to 5N or 5A exons in T98G cells.

| Minigene | Number of 5A clones | Number of 5N clones | Predominant splice variant |
| --- | --- | --- | --- |
| Wild-type | 8 | 2 | 5A |
| T35G | 1 | 7 | 5N |
| C46G | 2 | 8 | 5N |
| T35C+C46G | 2 | 8 | 5N |

In the second strategy, the products of PCR amplification were restriction digested with AvrII (cleavage site C|CTAG|G) (NEB, R0174S) which uniquely cleaves the amplification products at position 22 or products containing exon 5N, but not those containing exon 5A. The sequence of amplification fragments obtained before and after AvrII restriction digestion were verified by Sanger sequencing.

The results in HEK293 cells are compatible with increased inclusion of SCN8A exon 5N (and decreased inclusion of 5A) in mutants T35A, C46G, T35A+C46G, T51C and T78C compared with the wild type minigene, while increased inclusion of exon 5A was observed in mutants G45T, C46T and T78G, consistent with the predictions of high-throughput mutagenesis analyses (see below). Similar results were observed in T98G cells (Figure S2)

#### Transfection and analysis of indel library

Construction of the SCN8A Exon 5N (92 nt) 1K Indel Library

A library consisting of 1000 different sequence variants of exon 5N (SCN8A 5N exonic indel oligo library) was ordered as a single stranded Oligo pool to Twist Bioscience, with the following design:

-276 single substitutions (all possible single nucleotide substitutions at each exon position)

-276 deletions (all possible deletions of length 1,3,6 and 21)

-144 double (adjacent) substitutions in the windows of positions 32-38 and 43-49 (all possible adjacent substitutions within these regions).

-276 double adjacent substitutions outside of the windows of positions 32-38 and 43-49.

-9 double long-range substitutions involving the positions of the variants found in the patient (35,46).

-18 double long-range substitutions involving positions adjacent to the variants found in the patient (34,45) and (36,47).

-wildtype sequence.

The complete list of the 1,000 sequences, including their identifiers and nucleotide sequence is included in **Table** S2.

The sequence of the library corresponding to the wild type clone is 132 nucleotides long:

5’-aactttggtttgattctgcag

GTATATAACAGAGTTTGTAAACCTAGGCAATGTTTCAGCTCTACGCACTTTCAGGGTACTGAG GGCTTTGAAAACTATTTCGGTAATCCCAG

gtaagatggtccggggttg -3’

where non-capital letters indicate invariant flanking intronic sequences, while capital letters indicate wild-type exon 5N sequences.

Single stranded lyophilized oligo library (Twist Bioscience) was resuspended in 18 ul DNase-free water to obtain a concentration of 2 nanogram per µliter. 20 ng of resuspended single stranded oligo library were amplified with Accuprime Taq DNA polymerase system (Thermo fisher scientific, 12339016) for 12 amplification cycles and cloned in the pCMVBeta SCN8A e4-e5N-e5A-e6 vector by Gibson cloning using the following complementary primers:

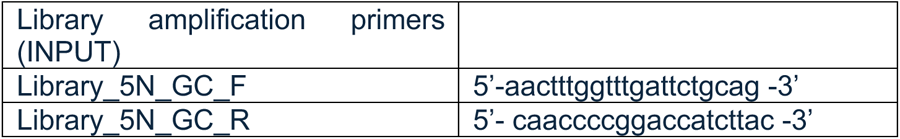

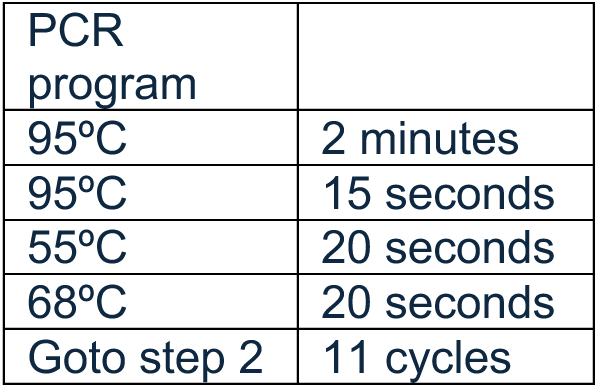
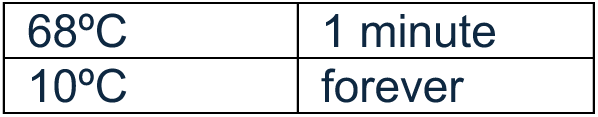

The products of 4 independent PCR reactions were combined and purified using the Quiaquick PCR purification kit (Quiagen, 50928106) eluted in 50 ml EB buffer and DNA concentration was measured using a Nanodrop instrument (dsDNA settings), obtaining a yield of 90,2 ng/ml.

#### Gibson DNA Assembly of SCN8A MB 1K library

The library was cloned in the pCMV SCN8A e4-e5A-e6 vector using 50 ng of amplified SCN8A_Indel_1K oligo library described above and 150 nanograms of linearized vector. pCMVbeta linearized vector, with no insert, was prepared by PCR amplification of the plasmid sequences distal to the location of library cloning using oligos complementary to flanking invariant sequences of the SCN8A oligo library:

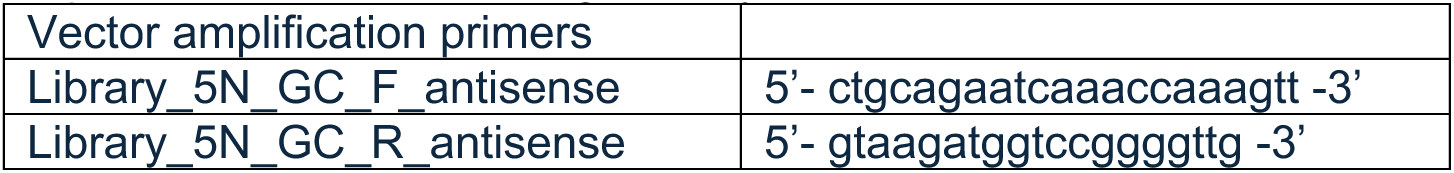

Amplified product was gel-purified (0.8% agarose) (Quiagen, 50928704). 5 µl of PCR product was phosphorylated, intramolecular ligated (circularization) and the original template removal by DpnI treatment using the KLD enzyme mix (NEB, M0554S) in a 10 µl reaction for 5 minutes at room temperature. Half of this reaction was transformed into competent cells, next day colonies were picked and verified by Sanger sequencing. Thus, the plasmid vector did not contain exon 5N sequences before cloning of the SCN8A 5N exonic library, to avoid any background of wild-type clones and full representation of the library.

For cloning the library, the vector: insert molar ratio was optimized to 1:8 to obtain a maximum number of cfu per transformation.

After Gibson assembly, the mix was transformed into competent Stellar competent cells (Takara, 636766), obtaining 979,200 cfu /ligation. Two independent ligations were set up, obtaining a total of 2,2 million clones. These were grown in LB media with Ampicillin for 18 hours and the cloned library was purified by DNA maxiprep (Qiagen, 50912163). 12 individual colonies were Sanger-sequenced to confirm the library design and diverse sequence representation.

#### Transfection conditions for generation of samples for Next Generation sequencing

100 nanograms of SCN8A 5N Indel library were transfected onto 180,000 -200,000 T98G cells in 6-well plates, in 6 biological replicates (SCNBR1-6). RNA was isolated 48 hours post-transfection, reverse-transcribed with vector-specific PT2 primer, and PCR amplified using the following primers corresponding to exonic sequences in SCN8A exons 4 and 6 that flank exon 5 (N or A) in the e4_e5_e6 inclusion products:

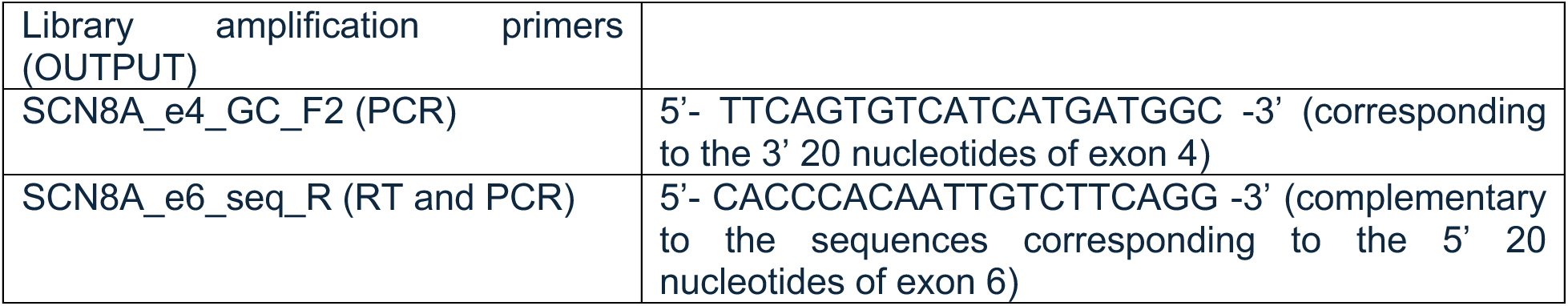

PCR amplification products corresponding to the single inclusion fragment band was excised from 6% acrylamide gels with the help of a clean scalpel and purified by “crush and soak” method (Molecular Cloning: A Laboratory Manual 4^th^ edition, 2012, Cold Spring Harnor Laboratory Press) and eluted in 200 µl acrylamide gel elution buffer with no SDS.

Therefore, the product of amplification corresponding to inclusion of the wild type 5N exon is 5’ TTCAGTGTCATCATGATGGC

GTATATAACAGAGTTTGTAAACCTAGGCAATGTTTCAGCTCTACGCACTTTCAGGGTACTGAG GGCTTTGAAAACTATTTCGGTAATCCCAG

CACCCACAATTGTCTTCAGGC-3’

To increase the complexity of the amplicons during Illumina sequencing, frameshifted forward and reverse primers containing NN- NNN- NNNN- or NNNNN-were utilized for PCR amplification. Thus, to generate INPUT samples (SCNTR1-3), a mix of the following eight frameshifted primers were utilized:

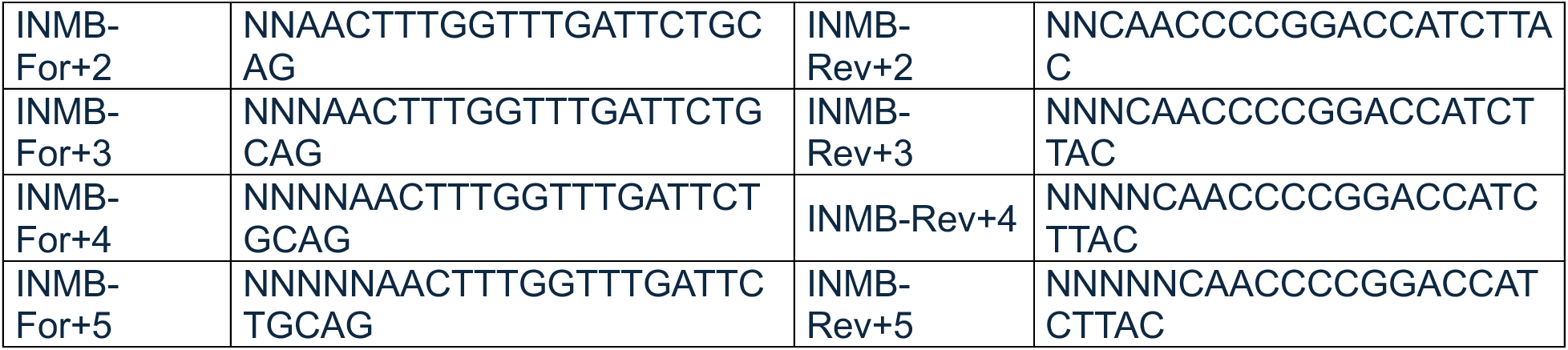

And for OUTPUT samples (SCNBR1-6), a mix of the following eight frameshifted primers were utilized:

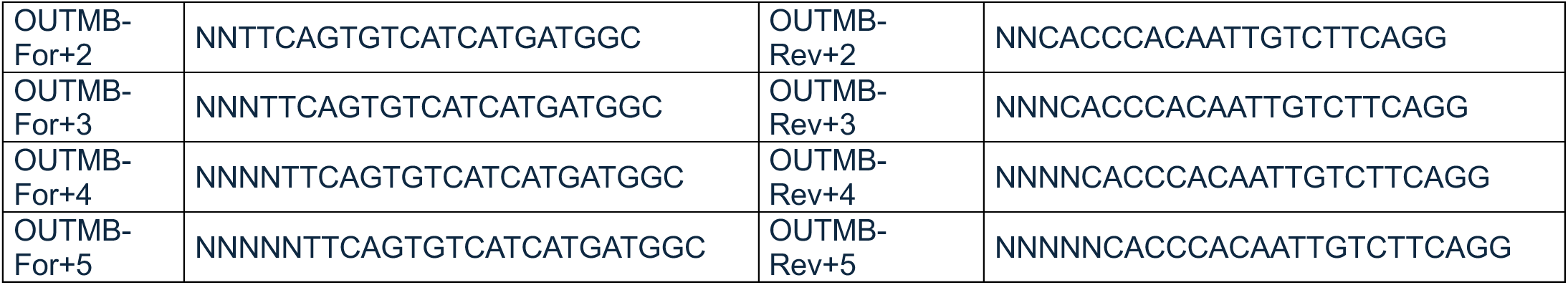

PCR amplification reaction of each replicate of INPUT or OUTPUT samples were prepared independently with a mix of eight frameshifted primers at 10 µM concentration.

PCR reaction was set up as follows

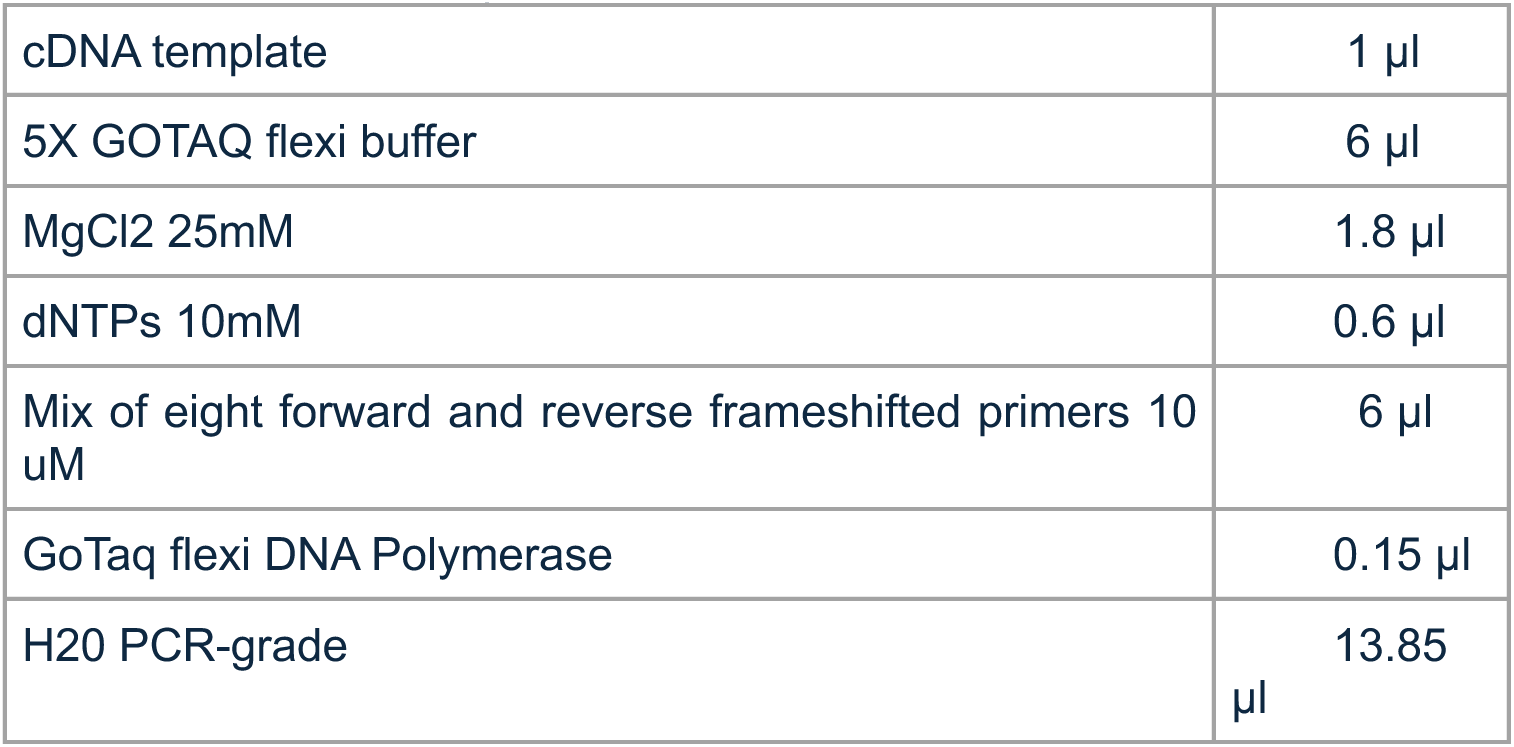

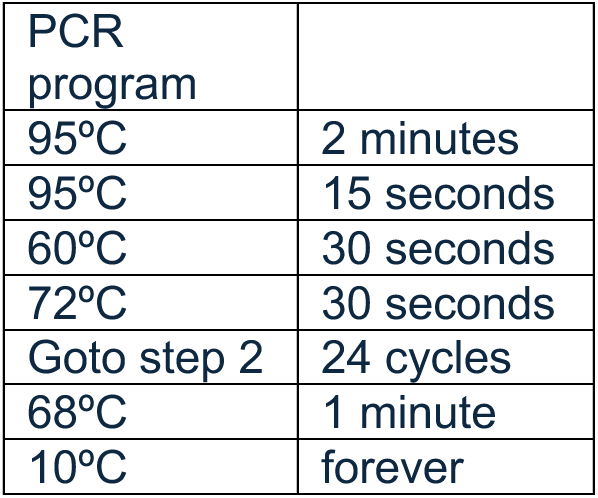

Full-length Illumina sequencing adapters were ligated using the NEBNext® Ultra™ II DNA Library Prep Kit from Illumina (NEB, E7645S) and samples were sequenced on a NextSeq2000 instrument in paired-end 2x150 bp mode.

#### Dimsum analysis of SCN8A exon 5N Indel library

Sequencing data were processed using ^18^. Briefly, DiMSum comprises an end-to-end pipeline for processing deep mutational scanning datasets from raw reads to measured sequences and their associated assay scores (plus errors). DiMSum was run with the following parameters:

cutadaptMinLength=“50”; cutadaptErrorRate=“0.5”;

vsearchMinQual=“30”; vsearchMaxee=“0.5”; startStage=“0”;

maxSubstitutions=“92”;

experimentDesignPairDuplicates=“TRUE”. Sequences with fewer than 100 reads were removed from the input sequencing experiment. Next, the inclusion estimates (PSI) of each variant were centered individually by subtracting from them the inclusion of the wildtype sequences. Inclusion values for each mutant are provided on Table S2.

Validation of inclusion heatmap using selected single nucleotide substitution clones and Avr II restriction enzyme digestion to distinguish between inclusion of exons 5N and 5A: the following mutants pCMV_SCN8A_e4_e5N_e5A_e6 wild-type, T35C, C46G, double mutant T35C+C46G, T51C, G45T, C46T, T98C, T98G were independently validated by transfection of individual mutants (or wild type) in the HEK293 and T98G glioblastoma cell lines (Figure S2a-e). The products of amplification were digested with Avr II (C|CTAG|G ; 1 site in exon 5N position 22) as described above.

Significance was determined by calculating the zstat_diff-- (inclusion_variant - inclusion_WT) / sqrt (error_variant^2 + error_WT^2) and performing a two-sided t-test over this z-statistic. The p-value for each substitution and deletion is shown in Table S2.

### Evolutionary analysis of Na^+^ and Ca^2+^ channel mutually exclusive exons

To assess conservation of mutually exclusive exon nucleotides we used positional conservation scores from the Zoonomia Consortium generated using 241 mammals. Bigwig files containing Zoonomia conservation scores were downloaded from: https://hgdownload.cse.ucsc.edu/goldenpath/hg38/cactus241way/cactus241way.phyloP.bw.^19^. Transcript, exon, and reading frame coordinates were obtained from the RefSeq version 40 and GENCODE version 38 human genome reference annotation releases. For all analyzed transcripts (Table S3) coding regions were split into MXEs and non-MXEs, then nucleotides were binned by position within a codon for that transcripts reading frame. The proportion of nucleotides conserved (Zoonomia conservation score ≥2.27) at each of the three codon positions was calculated for each category. To assess differences in conservation at codon position three between the MXEs and all other exons in the same gene we used these proportions as input for a beta binomial model. For each gene we independently assigned a uniform prior to the proportion of MXE nucleotides conserved:

P_MXE_ ∼ Beta(αʹ=1, βʹ=1)

Modelling the likelihood for proportion of conserved nucleotides as a binomial distribution:

k_MXE_ ∼ Binomial(N_MXE_, P_MXE_)

Giving us the posterior distribution:

P_MXE_|k_MXE_ ∼ Beta(α = k_MXE_ + αʹ, β= N_MXE_ - k_MXE_ + βʹ)

where:

P_MXE_ = Proportion of third position nucleotides conserved (≥2.27)

N_MXE_ = Total number of third position nucleotides

k_MXE_ = Observed number of conserved nucleotides (≥2.27)

We then separately modeled third codon position nucleotides for all other non-MXE exons in each gene using the same approach. For each transcript 10,000 samples were drawn from MXE and non-MXE posterior distributions and the log_2_ ratio of the posteriors was calculated as: log_2_(P_other_|k_other_) – log_2_(P_MXE_|k_MXE_). Points in plots represent the mean of the difference in posteriors with 94% credible intervals.

To assess sequence conservation in Ca^2+^ channel proteins full protein and MXE exon amino acid sequences were aligned using MUSCLE through EMBL-EBI webserver https://www.ebi.ac.uk/jdispatcher/msa/muscle?stype=protein ^20,21^. Alignments of full length protein isoforms were then used to generate a phylogenetic tree with PhyML 3.0 webserver using the default settings ^22^. Sodium channel exon and intron nucleotide sequence alignments and percent identity matrices were generating using the CLUSTAL Omenga webserver (https://www.ebi.ac.uk/jdispatcher/msa/clustalo?stype=protein).

### Antisense Oligonucleotides

ASOs 16 nucleotides in length were designed to exclusively target either *Scn8a* exon 5A or 5N using PFRED computational platform^23^. ASOs contain 2’MOE and phosphorothionate backbone throughout. Control ASO, ASO-CTL, was designed to not match to any transcript in the mouse genome and was previously used in *Scn8a* knockdown ASO studies^24^. ASOs used in vitro were manufactured by Integrated DNA Technologies (IDT). ASOs used in vivo were manufactured by Ionis Pharmaceuticals. The *in vivo* ASO-CTL was modified with a backbone developed by Ionis Pharmaceuticals.

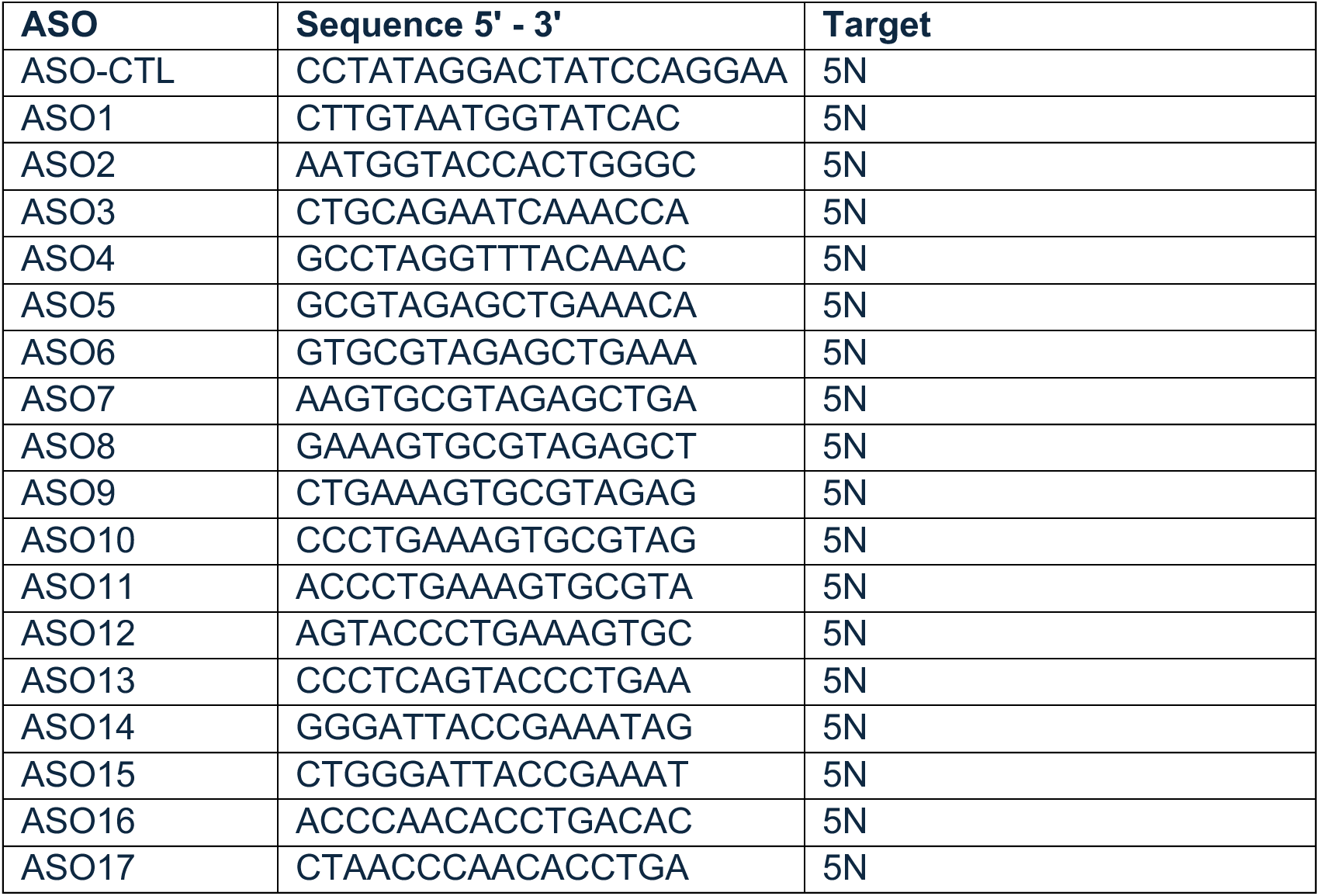
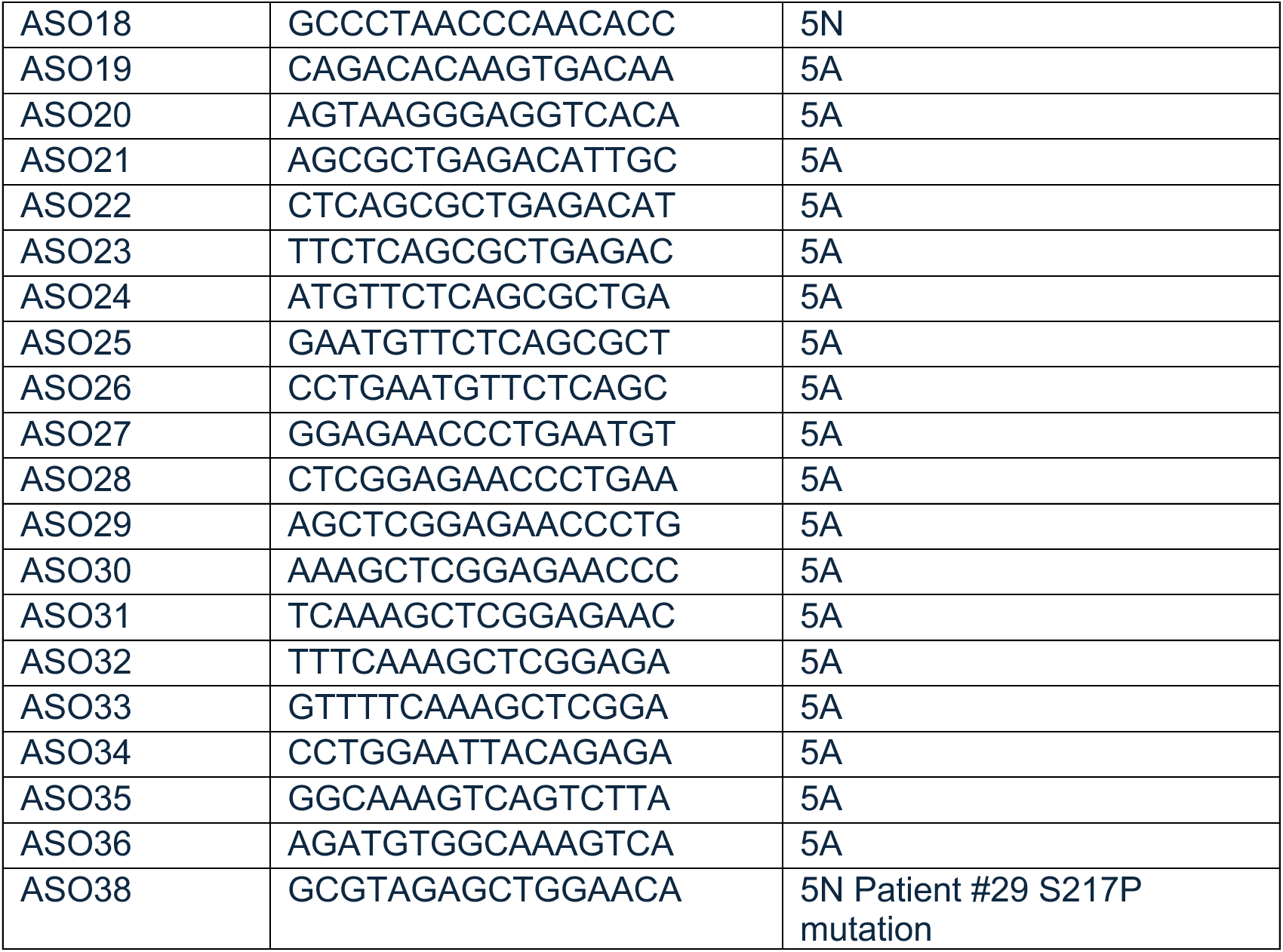

### Cell Culture

ND7/23 (92090903) and SH-SY5Y (94030304) cells were obtained from Millipore Sigma and cultured in DMEM with 10% FBS and 1% penicillin–streptomycin–glutamine (Fisher Scientific, 10-378-016). Cells were monitored for *Mycoplasma* contamination by PCR using the Universal *Mycoplasma* Detection Kit (30-1012 K; ATCC). Only *Mycoplasma*-negative cells were used in this study and all experiments were conducted within 15 passages of thawing. Cells were cultured at 37°C, and 5% CO_2_.

### Primary Cortical Neuron Culture and ASO Treatment

Cortexes were isolated from litters of C57Bl/6 mice at P0. All cortexes from the litter were pooled into one sample and cells were isolated, prepared and cultured for 7 days following previously published methods, in 24 well plates (Corning, 3524) ^25^. Cells were treated with 1000nM ASO via gynmotic transfection for 24hrs before RNA isolation.

### *In Vitro* ASO Treatment

ND7/23 and SH-SY5Y cells were seeding at 30% confluency (approximately 150,000 cells) in a 12-well plate (Corning, 3513). Cells were treated with 500nM ASO using Lipofectamine 3000 (ThermoFisher, L3000001) transfection following the manufactures protocol for 24hrs, before RNA isolation.

### RNA Analysis

Total RNA was extracted using Zymo RNA Preparation Kit (Fisher Scientific, 50-444-597). cDNA was synthesized from RNA using the High-Capacity cDNA Reverse Transcription Kit (ThermoFisher, 4368814). PCR amplification targeted *Scn1a*, *Scn2a*, *Scn3a* or *Scn8a* was completed with 1000ng of cDNA using Phusion Green High-Fidelity DNA Polymerase (ThermoFisher, F534S). PCR products were digested with either a 5A targeting enzyme, (AvaII, Thermofisher ER0311) or a 5N targeting enzyme (StyI, ThermoFisher ER0411). Digested products were separated a 3% agarose (Fisher Scientific, R0492) gel made with TBE (ThermoFisher, B52) and stained with a SYBR Safe DNA Gel Stain (ThermoFisher, S33102). Band intensity of digested and undigested bands were quantified using ImageJ (NIH). Percent 5A and 5N inclusion was calculated by averaging the ratio of digested and undigested band intensities for each exon targeting enzyme. Technical replicates were averaged to produce a single biological replicate. Assay validation was conducted using SCN8A exon 4-6 geneblocks (IDT) with inclusion or Exon 5A or 5N. Geneblocks were mixed in defined ratios prior to PCR amplification to determine precision and accuracy of the assay.

Primers used include:

*Scn1a* (exons 4-6) 5’- TTTACTTTCCTTCGGGATCCAT-3’ (forward) and 5’-ATACGCTCAGACAGAACACAG-3’ (reverse); *Scn2a* (exons 4-6) 5’-CCTGGAACTGGCTGGATTT -3’ (forward) and 5’- GGCCACTGCAAGCATTTATT -3’ (reverse); *Scn3a* (exons 4-6) 5’- CCATGGAACTGGCTGGATTT -3’ (forward) and 5’-GGCCACTGCAAGCATTTATTC -3’ (reverse); *Scn8a* (exons 4-6) 5’-TGGAACTGGTTAGATTTCAGTGT -3’ (forward) and 5’- CAACACACTTGTTTCGAAGGTT -3’ (reverse).

### qPCR

Total RNA was extracted using Zymo RNA Preparation Kit (Fisher Scientific, 50-444-597) from ND7/23 or SH-SY5Y cells treated with 500nM ASO via lipofectamine transfection. cDNA was synthesized from RNA using the High-Capacity cDNA Reverse Transcription Kit (ThermoFisher, 4368814). qPCR was conducted using 1000ng of cDNA with a TaqMan Gene Expression Master Mix (ThermoFisher, 4369016) and the following gene expression kits, *Scn8a* (ThermoFisher, Mm00488110_m1), *Gapdh* (ThermoFisher, Mm99999915_g1), SCN8A (ThermoFisher, Hs00274075_m1), GAPDH (ThermoFisher, Hs02786624_g1). GAPDH was used as an internal control. SCN8A gene expression was quantified using ΔΔ*C*_t_ method normalizing to the appropriate control.

### Induced Pluripotent Stem Cell (iPSC) generation

Fibroblasts were collected from the patient at Boston Children’s Hospital and prepared as previously published ^26,27^. Patient #29 has 3 known variants in SCN8A which are in cis: S217P, which is pathogenic, and impacts both the splicing and biophysical properties of Na_v_1.6, T144S, which was determined to be benign and not affect Na_v_1.6 biophysical properties and R220R, which only impacts splicing ^3^. Written consent was obtained from patients and sample collection was performed according to the institutional review board guidelines (IRB#: P00016119). Skin punch biopsies were rinsed with dPBS (ThermoFisher #14190250) and incubated at 37 °C in 0.5% Dispase (ThermoFisher #17105041) for 1 h. After one rinse with dPBS, epidermis was removed, each sample was sectioned into four quarters, and each quarter was placed in a 0.1% gelatin-coated well (Millipore #ES-006-B), weighed down by a coverslip on top. Samples were cultured at 37 °C in 5% CO_2_ in fibroblast media, consisting of 16% FBS (ThermoFisher #10439-024), 1x Penicillin/Streptomycin (ThermoFisher #15140-122), and 83ug/mL Primocin (InVivoGen #ant-pm-1). After about one month, all fibroblasts migrated out from each skin sample were dissociated with 1x Trypsin (ThermoFisher #15400054) and seeded into a 60 mm tissue culture dish. This is considered as passage 0 (p0) and mycoplasma testing (Lonza #LT07-418) was conducted for fibroblasts at this stage. Fibroblasts were further expanded in 100 mm dishes for two more passages, two dishes per passage. Fibroblasts in each 100 mm dish were cryopreserved into one cryovial in 10% DMSO (Sigma #D2650), 40% FBS, and 50% fibroblast media. Prior to reprogramming, one vial of fibroblasts at p2 was thawed into two 100 mm dishes and cultured in fibroblast media without primocin. Human induced Pluripotent Stem/Embryonic

Stem cell (iPS/ES) controls included two commercially available non-isogenic lines Kolf2.1J^28^ (iPSC, JAX JIPSC001000) and RUES2 (hES, WiCell RUES2). Newly generated pS217P iPSC lines were reprogrammed from fibroblasts using the CytoTune iPS 2.0 Sendai Kit (Thermo Fisher Scientific, A16517) following the manufacturer’s instructions. All cell lines were karyotyped to screen for spontaneous mutations prior to expansion and use. Both control lines were used to evaluate SCN8A Exon 5A:5N expression. The Kolf2.1J line was explicitly used as a reference for neuronal activity by multielectrode array. Pluripotent cells were cultured on Cultrex UltiMatrix (10ug/cm^2^, R&D systems, BME001) with Stemflex (Thermo Fisher Scientific, A3349401) media changes every 24-48 hours. Cells were passaged every 2-3 days at a 1:10 ratio using ReLeSR (STEMCELL Tech, 100-483) and 5uM Rock Inhibitor (Y-27632 dihydrochloride, Tocris, 1254). Long-term iPS/ES stocks were generated by resuspending cells in CryoStor CS10 (Millipore Sigma C2874) and cryopreserving according to manufacturer’s instructions. All pluripotent cell lines were routinely karyotyped for common abnormalities using the hPSC Genetic Analysis Kit (StemCell Tech, 07550) following manufacturer’s instructions. Cell cultures were tested for the presence of mycoplasma every month using the MycoAlert Mycoplasma Detection Kit (Lonza LT07-318) following the manufacturer’s protocol.

### iPSC-Neuron Generation and Differentiation

*Generation of Stable TetO-NGN2 iPSCs:* Pluripotent cells were seeded on Cultrex UltiMatrix (10ug/cm^2^, R&D systems, BME001) coated 24-well plates at 100k cells per well in Stemflex with 5uM Rock Inhibitor. The following day, cells were infected with pLVX-UbC-rtTA-Ngn2:2A:EGFP (Addgene 128288) lentivirus at MOI 10 in Stemflex. After 48 hours incubation 37 °C, 5% CO_2_, the transduced cells were selected with 1µg/ml Puromycin for an additional three days to generate a pure population of NGN2-iPSCs. Puromycin-resistant NGN2-iPSCs were expanded through two subculturing cycles into 6-well plates, then 10cm plates before generating long-term stocks.

#### Differentiation of inducible Cortical Neurons

On DIV 0, NGN2-iPSCs were seeded on Cultrex UltiMatrix (10ug/cm^2^, R&D systems, BME001) coated plates, in Stemflex with 2µg/ml Doxycycline Hyclate (ThermoFisher, J60579.14) and 5uM Rock Inhibitor. Media was refreshed to Stemflex with 2ug/ml doxycycline the following day. After 48h of induction onset, neurons were expanded 2-fold using Gentle Dissociation Reagent to Cultrex-coated plates. Media was changed to Cortical Neuron Media (CNM: 0.5x Neurobasal, 0.5x Brainphys, 2% Glutamax Supplement, 1x N2 Supplement, 1x B27 Supplement, 2ug/ml laminin, 500µM db-cAMP, 20ng/ml recombinant BDNF, 20ng/ml human GDNF, 10ng/ml human NT3) with 2ug/ml doxycycline and 5uM Rock Inhibitor. Media was refreshed daily with CNM supplemented with 2ug/ml doxycycline until DIV 4, when neurons were re-plated for the final time before experimentation. On DIV 4, neurons were dissociated with Gentle Cell Dissociation reagent and plated with CNM supplemented with 2ug/ml doxycycline, 5uM Rock Inhibitor, and 5uM Aphidicolin. Neuron media was fully refreshed to CNM on DIV6, removing doxycycline and aphidicolin. Neurons were maintained in CNM with half-media changes every 2-3 days until maturation on DIV 14 and subsequent experimentation.

#### Immunocytochemistry of iPSC-Neurons

Human iPSC-derived neuronal cultures for immunofluorescent staining were generated according to the same protocol described in iPSC-Neuron Generation and Culture. On DIV 4, neurons were dissociated into a single-cell suspension (Accutase, Stemcell Technology, 07922) and seeded at 25,000 cells per well, on a cultrex-coated ibidi 18-well µ-slide (ibidi, 81816). Cells were cultured for an additional 10 days to reach maturity before fixation, with half-volume neuronal media changes every two to three days as required. Neurons were fixed with 4% para-formaldehyde (PFA, Life Technologies 28908) for 30 minutes at room temperature, then washed three times with 1xPBS before blocking/permeabilizing with SuperBlock Blocking Buffer (ThermoFisher, 37515) for one hour at room temperature. Neurons were then incubated with primary antibodies overnight at 4°C in SuperBlock. Primary antibodies used were: anti-SCN8A (1:250, Invitorgen, PA5-37276), anti-MAP2 (1:500, Invitrogen, PA1-10005). The following day, neurons were washed three times with SuperBlock to remove primary antibody, then incubated with secondary antibodies overnight at 4°C in SuperBlock. Secondary antibodies used were: goat anti-chicken 488 (1:1000, ThermoFisher, A-11008), goat anti-rabbit 546 (1:1000, ThermoFisher, A-11035). Neurons were then washed once with 1xPBS and incubated for 5 minutes with 1ug/ml DAPI (ThermoFisher, 62248). Neurons were washed an additional three times to remove DAPI, and mounted in ibidi Mounting Medium (ibidi, 50001) before imaging on an Andor Dragonfly 200 Spinning Disk Confocal microscope (Fusion software version 2.4; 20x objective).

#### ASO Treatment and Isoform Expression Analysis

iPSC-Neurons were cultured and differentiated to DIV 14 as described above. On DIV 14, neurons were fed by half-media refresh 30 minutes prior to treatment with 500nM ASO using Lipofectamine RNAiMAX Transfection Reagent (ThermoFisher, 13778075) per manufacturer’s instructions. Neurons were incubated at 37°C with 5% CO^2^ for 72 hours. On DIV 17, ASO concentration was diluted by a half-media change to the existing ASO-treated neurons. RNA was harvested on DIV18 five days post-ASO treatment using TRIzol (ThermoFisher, 15596026) and immediately extracted using the Direct-Zol RNA miniprep kit (Zymo, R2050). RNA was stored at -80°C until further experimentation.

#### Microelectrode Array (MEA) Cell Culture

Human iPSC-derived neuronal cultures for Multi-Electrode Array recordings were generated according to the same protocol described in iPSC-Neuron Generation and Culture. On DIV 4, neurons were dissociated into a single-cell suspension (Accutase, Stemcell Technology, 07922) and plated in a 50µl droplet of 200,000 cells per well, on a pre-coated Cytoview MEA 48 plate (Axion Biosystems, m768-tMEA-48B). Pre-coating consisted of placing a droplet of 50ul Cultrex Ultimatrix RGF BME (diluted to 80ug/ml, R&D Systems, BME001) directly over the electrode grid, incubating at 37°C for at least 30min. Immediately after plating cells, sterile water was added to the plate reservoir, and the MEA plate was incubated at 37 °C, 5% CO^2^ in a cell culture incubator. After two hours of incubation to allow the cells to adhere, 150 µl additional neuronal medium was added to each well. Cells were cultured for an additional 10 days to reach maturity before beginning recording, with half-volume neuronal media changes every two to three days as required.

*ASO Treatment on MEA:* Spontaneous baseline neuronal activity was recorded on DIV 14 prior to ASO treatment to confirm active electrodes and neuronal health. After baseline recording, neuronal media was refreshed to 200µl containing 500nM ASOs administered by free-uptake and incubated at 37°C with 5% CO^2^ for 72 hours. On DIV 17, ASO concentration was diluted by a half-media change to the existing ASO-treated neurons. Spontaneous neuronal activity was measured again at five days post-ASO treatment.

#### MEA Recording

MEA recordings were taken by a Maestro Classic MEA system (Axion Biosystems) using Axion Integrated Studio (AxIS) Acquisition software (version 2.1.5). Neuronal media was refreshed 24h prior to recording, normalizing metabolic conditions of all neurons. Cultures were maintained at 37°C with 5% CO_2_ during the recording session. Plates were equilibrated in the system for 15 minutes to prevent mechanical or environmental artifacts. Neuronal activity was acquired at a sampling frequency of 12.5 kHz, using a Butterworth band-pass filter (200-3000 Hz) and adaptive spike detector set at 5.5 standard deviations from mean background noise. An electrode was defined as active with a mean firing rate greater than 0.017 Hz, wells with ≤30% active electrodes were not included in data analysis. Data acquisition for each condition consisted of a minimum of three 5-min recording sessions over the course of 30min. Thresholded spike data, weighted mean firing rate, bursting frequency, and bursting duration were analyzed using Axion NeuralMetricTool software (version 2.1.5). A bursting event (burst) was defined as five or more spikes in a row, with a maximum interspike interval of 100ms, using the same analysis software.

### Animals

All animal protocols were approved by the Tufts Institutional Animal Care and Use Committee. Experiments used C57/Bl6 and S217P^+/-^ mice which were developed by GeneOway, modeled off of Patient #29’s mutations. Patient #29 has 3 known variants in SCN8A which are in cis: S217P, which is pathogenic, impacts splicing and biophysical properties of Nav1.6, T144S, which was determined to be benign and not affect Nav1.6 biophysical properties and R220R, which only impacts splicing. The mouse model is referred to as S217P^+/-^. Breeding S217P^+/-^ mice were kept on 8mg/kg diet of sodium channel inhibitor GS967 (Selleckchem, S5274). Mice were returned to normal diet upon weaning. Due to lack of colony viability equal numbers of male and female mice were not used in experiments. Mice were housed under 12/12-h light/dark cycles at approximately 21 °C and 30–70% humidity.

### ASO neurotoxicity studies

Intracerebral ASO delivery was performed as previously published^29,30^, with 700ug of ASO injected. The acute neuronal activation scoring system was recently developed to evaluate neurotoxicity, focusing on capturing neuronal activation vs. sedation ^31^ ^32^. Mice are given a score from 0-7, with 0 represents animals alert but at rest or performing maintenance activities such as sleeping, eating, grooming or nesting. A score of 1 was assigned if animals had hunched backs with signs of pain or were freezing in the cage. A score of 2 was assigned when shivering or spontaneous shaking occurred. A score of 3 represented animals with stiff and raised tails and/or hyperactive scuttling around the cage. A score of 4 was used when severe limb stiffness that impaired forward movement was present (Stiff limbs when walking). A score of 5 was given if they presented evident popcorning (vigorous jumping and hyperexcitability) or hopping behavior. A score of 6 represented any kind of seizure. A score of 7 was given if the animal died during the 2hr evaluation. Mice were evaluated at 3hrs and 24hrs post-ASO injection. For inflammatory marker gene expression analysis, RNA was extracted using TRIzol® (TRIzol Reagent, Invitrogen, cat # 15596026). Then, total RNA was isolated using the RNeasy 96 Kit (Qiagen, Germantown, MD). RT-qPCR was performed using the EXPRESS One-Step Super-Script qRT-PCR kit. Gene-specific primers were used from^31^ ^32^ and purchased from IDT technologies (Coralville, IA). For ASO-treated animals, the expression level of *Gfap*, *Aif1*, and *ATXN3* genes were normalized against *Gapdh* and this was further normalized to the mean level in vehicle-treated animals.

### *In Vivo* ASO Treatment

At postnatal day 1 (P1) mice were anesthetized using hypothermia. Mice were placed on a piece of plastic film on top of ice. Once anesthetized, they receiving 1uL of 30ug/uL ASO diluted in PBS to both the left and right cerebral ventricle for a final dose of 60ug ASO. ASO was injected using a glass filament (Drummond, 3-000-203 G/X) pulled on a P97 Stutter Micropipette Puller, at Heat = 660, Pull = 35 and Velocity = 35. Coordinates for injections were as follows: 1mm mediolateral and 2mm superior to lambda. Injections took less than 2 minutes, mice were placed on heating a pad before returning to the dam.

### Mouse Pup Behavior

#### Negative Geotaxis

Mouse pups were assed daily from P7-P14 using negative geotaxis. Pups were placed face downward on a ramp at an 80° angle. The latency to turn 180° degrees to face upward was recorded. Pups that did not complete the task within 30 seconds were considered to have failed and returned to the dam.

#### Righting reflex

Righting reflex was assed daily from P5-P14 for ability to right. Mice were placed upside down on a clean, padded surface. The time for mice to right themselves to an upward position was recorded. Mice were considered to have failed the task if exhibiting a full body tension, a sign of distress or if failed to right itself within 30 seconds, at which point the mice were returned to the dam.

### Pup video recording for movement phenotype and seizure occurrence

Mice pups (P14-P20) were recorded daily in a divided clear plastic container for 5-10 minutes. Videos were scored for seizure occurrence using a previously described Racine scale for pups^33–35^. Number of seizures and falls were counted and adjusted to represent occurrences per 10 minutes.

Videos were independently scored by two experimenters and compared for consistency. Experimenters were blinded to genotype and ASO treatment.

### Adult Animal Behavior

Behavioral testing of adult mice (6-10 weeks of age) occurred in the Tufts Circuits and Behavior Core. Adult behavior was conducted on individually identifiable tattooed mice. Mice were handled by the experimenter prior to behavior tests for habituation. Apparatuses were cleaned with 70% ethanol before and between animal trials. Behavioral tests were conducted between 10.00-13.00 hr, with male mice trialed before female mice. The same cohorts of mice completed open field, y-maze and light dark box.

#### Open Field

Mice were individually recorded in 36cm x 36cm opaque open field arena for 10 minutes. Behavior was tracked with EthoVision software (Noldus Information Technology). Ethovision was used to calculate total distance traveled, average velocity, and time spent in the center and borders of the arena. Distance and velocity were used to determine locomotor activity, while time spent in center versus borders was an indicator of an anxiety-like behavior.

#### Y-Maze

A three-armed apparatus was used for the y-maze. Each arm was 42cm in length. Mice were recorded in the Y-maze for 10 minutes and monitored using EthoVision software (Noldus Information Technology). EthoVision calculated total distance traveled, average velocity, number of entries into each arm, and time spent in each arm. Spontaneous alternations were calculated and used as a measure of spatial working memory of the mice.

#### Light/Dark Box

Mice were placed in a two-chamber light/dark box with equal size compartments. Mice were tracked in the light/dark box over 10 minutes using a 22cm x 43cm photobeam frame, with MotorMonitor software (Hamilton-Kinder). MotorMonitor recorded time in either the light or dark compartments and entries into each compartment. On day one trials were conducted at ambient light conditions (25lux), and on day two at bright light conditions (500lux). MotorMonitor software then analyzed the recorded for outputs of Time in Dark, Time in Light, Dark Entries and Light Entries.

### Adult video Recording

F1 S217P^+/-^ and WT mice (7-8 weeks old) were individually placed in an 38cm x 28cm open arena with pellet bedding, material bedding, food and water. Mice were recorded for 1hr in the arena. Videos were analyzed for time active (moving, grooming, eating, and drinking) and sedentary as well for vertical ambulation by climbing.

#### Adult, chronic ECoG *in vivo* recordings and seizure burden quantification

##### Surgery

ECoG electrode-surgery was performed on adult 7–9-week-old F1 S217P^+/-^ and WT mice as previously described^36^. In brief, mice were anesthetized with isoflurane (3% v/v for induction, 1.5% v/v for maintenance, 2 l/min O_2_ flow rate). Systemic analgesia with <u>buprenorphine</u> (0.1 mg/kg, SC) and additional local analgesia around the incision site with <u>bupivacaine</u> (4 mg/kg, SC) were provided for all mice. Mice were then placed in a stereotactic frame, dorsal side of skull shaved and disinfected. A scalp incision was made, the skull surface cleaned. Four burr holes were drilled without puncturing the dura mater (Drill bit diameter 0.7 mm, Item No. 19007-07, Fine Science Tools, Foster City, CA, USA). Anterior burr holes carried ECoG screw electrodes for each hemisphere. Posterior burr holes accommodated ground and reference electrodes. Stereotaxic coordinates of the four drill holes: Anterior burr holes at −0.6 mm (bregma, AP axis) and +/-2 mm of midline (ML), posterior burr holes at –2.6 mm (AP) and +/-2.5 mm (ML). Four stainless steel screw electrodes with attached silver wires (# 8403, Pinnacle Technologies, Wyckoff, NJ, USA) were screwed in and fixed with dental cement. Silver wires were soldered to a headmount (# 8402, Pinnacle Technologies, Wyckoff, NJ, USA) and further secured with dental cement. Mice recovered for at least seven days before chronic ECoG -recordings start.

##### Recordings

24/7 ECoG recording signals were acquired using a preamplifier (100 x gain, 1 Hz high-pass filter, # 8213, Pinnacle Technologies, Wyckoff, NJ, USA) and recorded using custom-made scripts running LabChart Pro software (AD instruments, Colorado Springs, CO, USA) with a 1 kHz sampling rate and 300 Hz low pass filter. Raw data was pre-processed (100 Hz low pass 8^th^ order Butterworth filter) and inspected for recording artifacts. Seizures were detected by a semiautomated procedure confirmed by manual review of the output and original ECoG traces by an experienced, blinded investigator. Paroxysmal events with clear tonic and clonic phases lasting for at least 10 seconds were counted as seizures. Seizure incidence and duration were used to determine seizure burden in mice.

### Brain immunohistochemistry

#### Immunofluorescence

Brains dissected from mice <P30 were fixed in 4% paraformaldehyde. Brains of mice >P30 were prepared by transcardial perfusion with PBS and fixed in 4% paraformaldehyde. Fixed brains were sectioned at 40μm using a ThermoFisher Microm HM 525 cryostat. Brain sections were blocked with blocking buffer (5% normal goat serum and 1% bovine serum albumin in PBS) for 1hr at room temperature. Adult mouse brains were stained with NEUN (1:500, Millipore, Mab317) and PV (1:1000, Abcam, ab256811), diluted in PBS with 0.2% Triton X-100 and 5% blocking buffer. Young mouse brains were stained with NEUN (1:500, Millipore, Mab317), PV (1:1000, Synaptic Systems, 195308) and ASO (1:10,000, Ionis Pharmaceuticals) diluted in PBS with 0.2% Triton X-100 and 5% blocking buffer. Cortical sections were incubated with diluted primary antibodies overnight at 4 °C. Secondary antibodies (goat anti-rabbit 647(Jackson ImmunoResearch Labratories), goat anti-mouse 555(Jackson ImmunoResearch Labratories) and goat anti-guinea pig 488 (Invitrogen)) were diluted 1:500 in PBS with 5% blocking buffer and added to brain sections for 2 h at room temperature. Slices were imaged using a 20x objective with a Keyence BZ-X710 microscope (F1 S217P^+/-^ and adult WT brains) or using a 20x objective using a Leica Thunder microscope (F2 ASO treated surviving adults and WT littermates, young F2 ASO-CTL and ASO-13 treated S217P^+/-^ and young ASO-CTL and ASO-13 WT mice).

#### Image analysis

Adult mouse brains isocortex regions of interest (ROIs) were determined using Aligning Big Brains & Atlases (ABBA), with Allen Brain Atlas V3p1 used^37^. Parvalbumin and NeuN positive cells were counted using ImageJ software. Isocortex area was calculated using ImageJ software and cells counts were then divided by area to determine cells/mm^2.^ Isocortex of young (P19) brains was determined by hand referring to the Allen Developing Mouse Brain

Atlas^38^. Parvalbumin cells were hand counted by two blinded researchers in ROIs, with counts between researchers averaged. NeuN cell counts and isocortex area were determined using ImageJ software. Cells counts were then divided by area to determine cells/mm^2^. ASO intensity in ASO13 treated animals was calculated by subtracting intensity of untreated animals, matching genotypes and brain regions.

### Statistical Analysis

Statistical analysis and visualization were performed using GraphPad Prism v8.4.3 and R studio unless indicated otherwise. Patient data was compared with the hypergeometric test, (* p<0.05 and *** p<0.001).

For the in vitro ASO screen, comparisons were conducted using one-way ANOVA with Dunnett’s multiple comparison test to untreated control (ns p>0.05, *p<0.05. ** p<0.01, *** p<0.001, **** p<0.0001). Behavioral assessments of WT ASO treatment were analyzed as follows: weight and negative geotaxis, two-way ANOVA with Tukey’s multiple comparison test comparing all groups and adult behavior, one-way ANOVA with Dunnett’s multiple comparison test to untreated control.

In vitro IPSC-derived comparisons were completed with Student’s t-test was performed when data were normally distributed (Kolmogorov–Smirnov normality test), and Mann–Whitney U was performed when data were not normally distributed all relative to untreated control (ns p>0.05, *p ≤ 0.05, **p ≤ 0.01, ***p ≤ 0.001, ****p≤0.0001).

In vivo characterization of F1 generation was analyzed as follows between WT and S217P^+/-^ mice: time active, climbs and *Scn8a* 5A:5N, multiple unpaired t tests; PV and NeuN immunostaining linear mixed modeling, comparing genotype and treatment as fixed effects and slices are considered technical replicates to biological animal (ns P>0.05, * P≤0.05, ** P ≤0.01, *** P≤0.001, ****P≤0.0001).

In vivo characterization of F2 generation was analyzed as follows: weight between WT and S217P^+/-^ mice by multiple unpaired t tests; seizure occurrence and falls by one sample t and Wilcoxon test (theoretical mean set to 0); and *Scn8a* 5A:5N inclusion by one-way ANOVA with Dunnett’s multiple comparison test to WT P0 (ns P>0.05, * P≤0.05, ** P ≤0.01, *** P≤0.001, ****P≤0.0001).

For ASO treated S217P^+/-^ and WT mice, survival was assed using the Mantel – Cox test between S217P^+/-^ ASOCTL and ASO13. Seizures and falls were analyzed by unpaired *t*-tests. *Scn8a* 5A:5N was analyzed by one-way ANOVA with Dunnett’s multiple comparison test to S217P^+/-^ ASOCTL. Time on back and side was analyzed by Kruskal-Wallis test with Dunn’s multiple comparison to ASOCTL. PV and NeuN immunostaining were analyzed using linear mixed modeling, comparing genotype and treatment as fixed effects and slices are considered technical replicates to biological animal. ASO immunostaining was analyzed using linear mixed modeling, comparing genotype and brain region as fixed effects and slices are considered technical replicates to biological animal, (ns P>0.05, * P≤0.05, ** P ≤0.01, *** P≤0.001, ****P≤0.0001).

## Acknowledgements

We acknowledge the help of Elizabeth Buttermore and Mustafa Sahin from Boston Children’s Hospital for the patient fibroblast culture for iPSC generation, the team at Genoway for generating the S217P^+/-^ mouse model, the Oudin family for supporting the generation of the mouse model, and all our friends and family who donated to the SCN8A Epilepsy Research Fund at Tufts University to support this work. We also acknowledge all the SCN8A families and patients who have shared their data and experiences in the SCN8A registry, especially our daughter Margot, who motivated us to start this work.

## Author Contributions

H.B.D., C.B.B., C.D. and M.J.O. conceived the project and designed experiments. H.B.D. designed and tested ASOs, developed 5A:5N splicing assay, tested ASOs in WT mice, conducted WT ASO treated behavioral assays, characterized S217P^+/-^ mice by survival, weight, neonatal behavior, video recording, EEG quantification, and immunofluorescence. H.B.D. treated S217P^+/-^ mice with ASOs and characterized them with above methods. S.A. generated the S217P^+/-^ iPSCs and H.K. differentiated and characterized S217P^+/-^ iPSCs, screened ASOs and generated MEA recordings with untreated and ASO-treated differentiated neurons. R.S. aided in iPSC-neuron culture and characterization B.M. and J.V. generated mini-gene and completed saturation mutagenesis of SCN8A and B.L and MT did analysis. Z.K. completed EEG cortical implants and analysis. J.B.F aided in ASO delivery and characterization of ASO treated S217P^+/-^ mice by immunofluorescence. D.M. and C.B.B. completed sequence conservation and splicing regulation analysis. V.R. aided in mouse care and neonatal behavior. C.Y. isolated fibroblasts and helped conceive original clinical ASO approach. N.T. aided in ASO treated WT mice behavioral assays. M.W. aided in ASO delivery and ASO treated S217P^+/-^ characterization with video recording and neonatal behavior. C.Z. aided in 5A:5N assay quantification. C.F.B. completed toxicology screening of ASOs in adult mice and provided ASO manufacturing for *in vivo* experiments. M.J.O, with M.H., analyzed exon 5 patient data. H.B.D. and M.J.O. with help from C.B.B. and H.K. wrote the manuscript with input from all authors. M.J.O. supervised all aspects of the work. All authors read the manuscript and provided feedback.

