## Supplemental data for "A splice-switching antisense oligonucleotide approach for pediatric genetic epilepsies"

### Supplemental Figure Legends

**Table S1: Patients with mutations in SCN8A Exon 5**

Table summarizing 52 patients with SCN8A mutations in exon 5 collected from the International SCN8A registry and published papers. Table shows mutation in DNA and corresponding amino acid change, sex, location of mutation, age of seizure onset, seizure types, where patient has seizure control and development.

**Table S2:** Sequences for mini-gene saturation mutagenesis and deletion analysis

**Table S3:** Table including the CDS for the transcripts used as well as hg38 coordinates for all the analyzed MXEs.

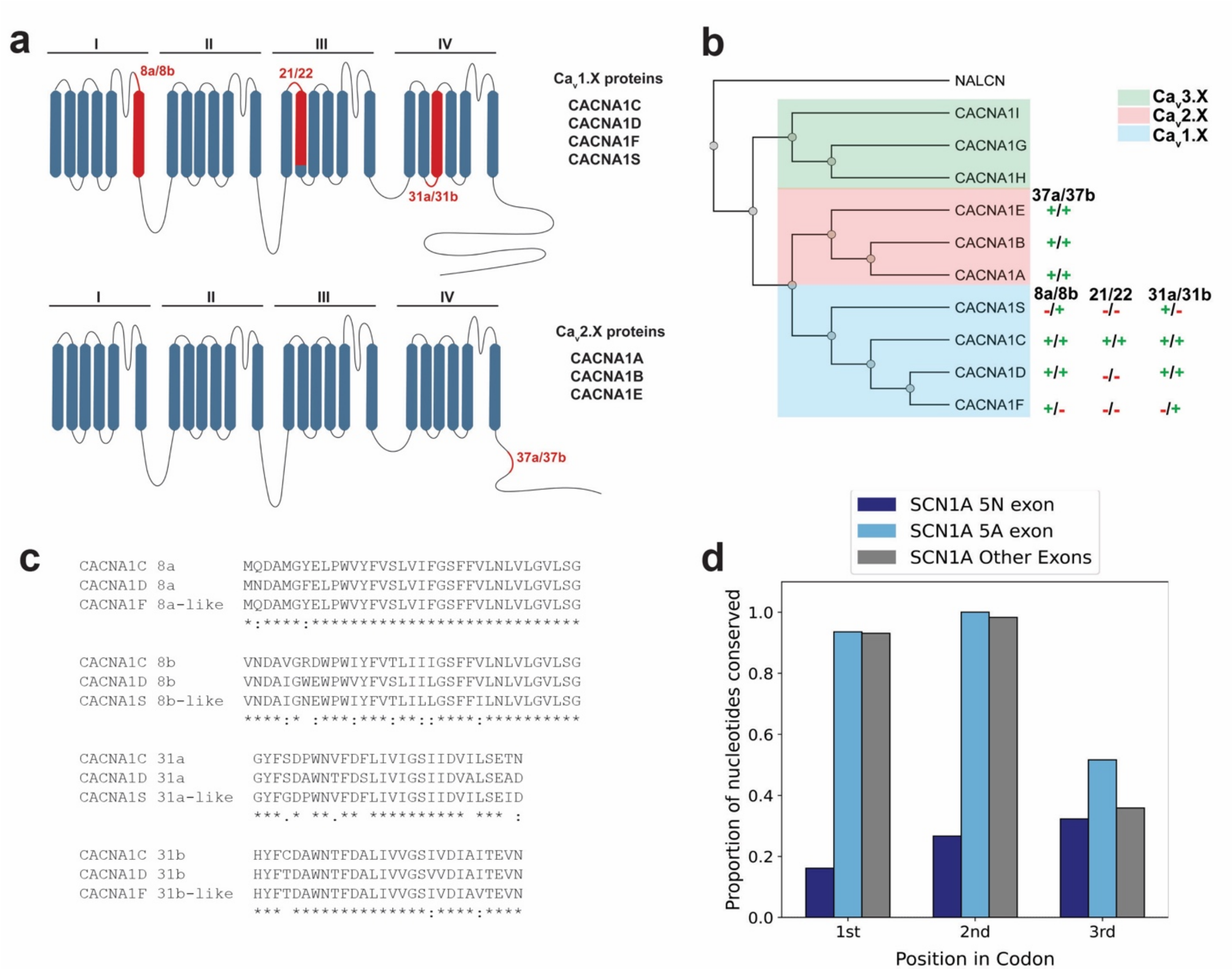


**Figure S1: Conservation of codon position three nucleotides in mutually exclusive exons in human ion channel genes. a)** Cartoon schematic of the structure of two families of Ca2+ channel proteins Ca$_v$1.X (top) and Ca$_v$2.X (bottom). Regions of the protein derived from mutually exclusive exons are depicted in red and labeled with the two exons names (upstream exon/downstream exon). Members of each protein family are listed to the right. Each of the four transmembrane domains are labeled above (I-IV). **b)** Phylogenetic tree depicting relationship of human $Ca^{2+}$ channel proteins with NALCN out group. To the right of the tree is a list of all known MXE pairs from each family. For individual proteins the presence or absence exon in the transcript is indicated as + and - respectively. **c)** Multiple sequence alignments of the MXEs 8a/8b and 31a/31b from CACNA1C and CACNA1D with the singleton exon equivalents found in CACNA1F and CACNA1S. **d)** Bar plot depicting the proportion of conserved nucleotides in SCN1A found in exons 5N, 5A, and all other SCN1A exons. Nucleotides are binned by position with codons.

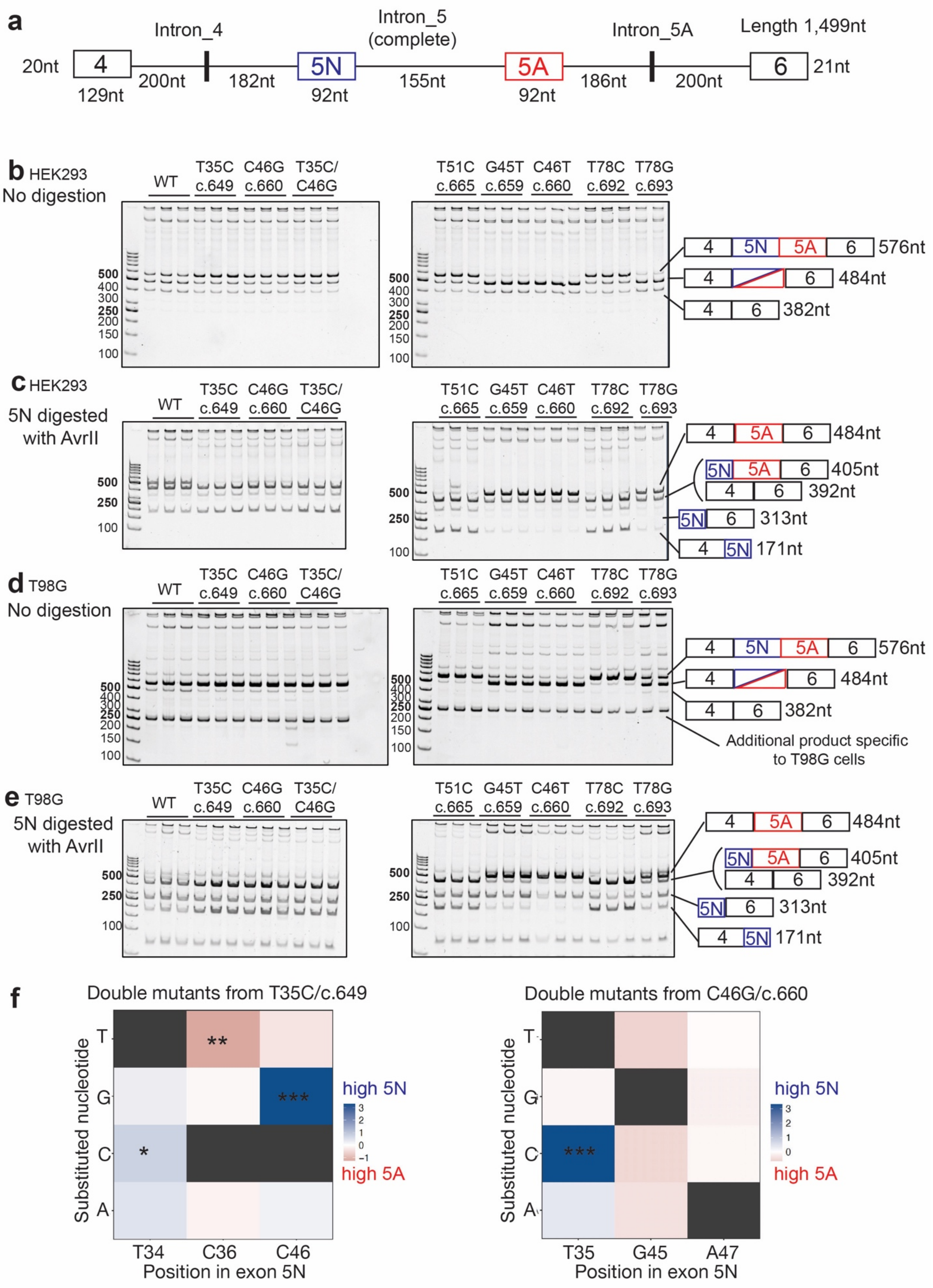
a
Intron_4
Intron_5 (complete)
Intron_5A
Length 1,499nt
20nt
4
129nt
200nt
182nt
5N
92nt
155nt
5A
92nt
186nt
200nt
6
21nt
b HEK293
No digestion
WT
T35C c.649
C46G c.660
T35C/ C46G
T51C c.665
G45T c.659
C46T c.660
T78C c.692
T78G c.693
576nt
484nt
382nt
c HEK293
5N digested with AvrII
484nt
405nt
392nt
313nt
171nt
d T98G
No digestion
Additional product specific to T98G cells
e T98G
5N digested with AvrII
f
Double mutants from T35C/c.649
Double mutants from C46G/c.660
Substituted nucleotide
Position in exon 5N
T34
C36
C46
T35
G45
A47
high 5N
high 5A

**Figure S2: Validation of saturation mutagenesis for single and double mutants**

**a)** Schematic of the mini-gene reporter containing SCN8A exon 4-5N-5A-6. The minigene spanned sequences from the 5' end of exon 4 to the 3' end of exon 6 cloned in an expression vector under a CMV promoter, with internal deletions of introns 4 and 5A due to their lengths. Validation by RT-PCR of data from the saturated mutagenesis analysis using the SCN8A mini-gene reporter, testing out 7 mutations in exon 5N which were predicted to have effects by the analysis of deep mutagenesis data: T35C or c.649, C46G or c.660, T51C or c.665, G45T or c.659, C46T or c.660, T78C or c.692 and T78G or c.693. Data shown for HEK293 cells with gels showing the PCR product (**b**) and after digestion with AvrII which cuts Exon 5N only (**c**). Data shown for T98G cells with gels showing the PCR products **d**) and after digestion with AvrII which cuts Exon 5N only (**e**). **f**) Quantification of effect of exon 5N inclusion changes for combinations of double nucleotide substitutions in SCN8A exon 5N. Black indicates wild type nucleotide at that position

**a**

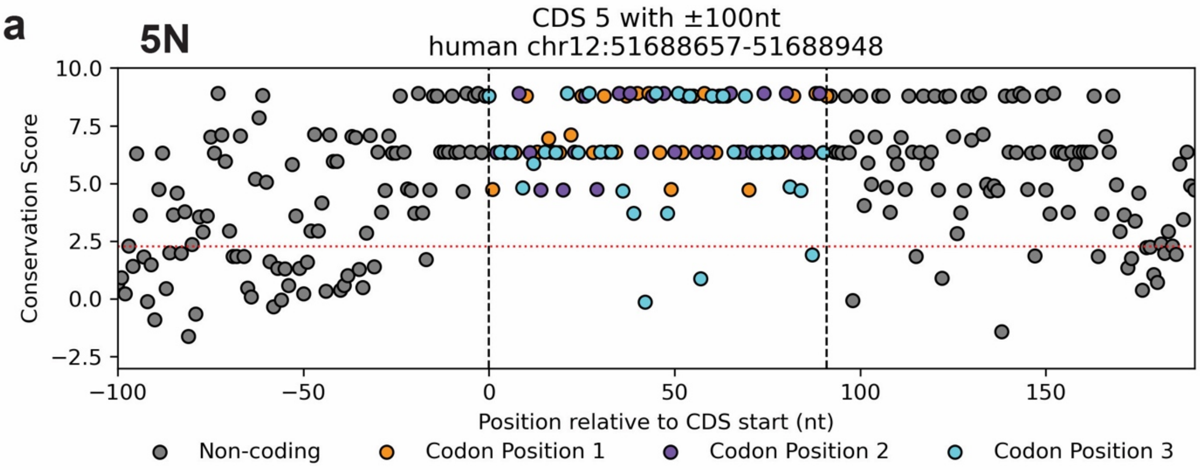


| SCN8A 5N nucleotide conservation | | | |
|---|---|---|---|
| Position category | Conserved nucleotides | Total nucleotides | Conserved propotion |
| Upstream intron | 65 | 100 | 0.65 |
| Downstream intron | 85 | 100 | 0.85 |
| Exon: 3rd position | 28 | 31 | 0.903 |

**b**

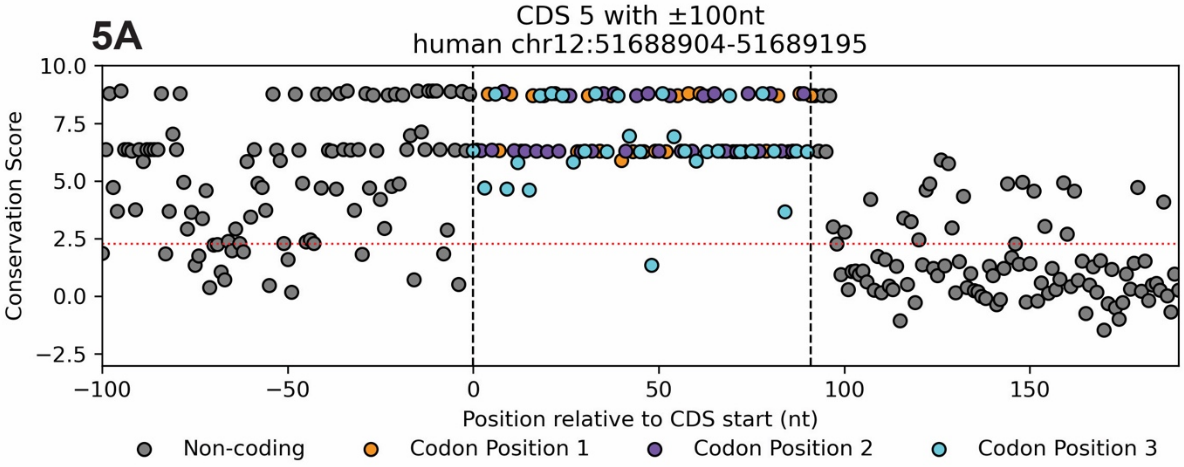


| SCN8A 5A nucleotide conservation | | | |
|---|---|---|---|
| Position category | Conserved nucleotides | Total nucleotides | Conserved propotion |
| Upstream intron | 82 | 100 | 0.82 |
| Downstream intron | 27 | 100 | 0.27 |
| Exon: 3rd position | 30 | 31 | 0.968 |

**Figure S3: Individual nucleotide conservation scores** for SCN8A 5N **a)** and 5A **b)** as well as 100 nucleotides upstream and downstream of each exon**.** X-axis shows the position of each nucleotide relative to the first nucleotide of the exon. The dotted red line indicates the 2.27 cut off to be considered conserved by Zoonomia. hg38 reference genome annotation position for the depicted region of SCN8A are indicated above each plot. Below each plot is a table indicating the number and fraction of nucleotides considered to be conserved (i.e. meeting the 2.27 cut off) for each of the three regions (exon and two flanking introns).

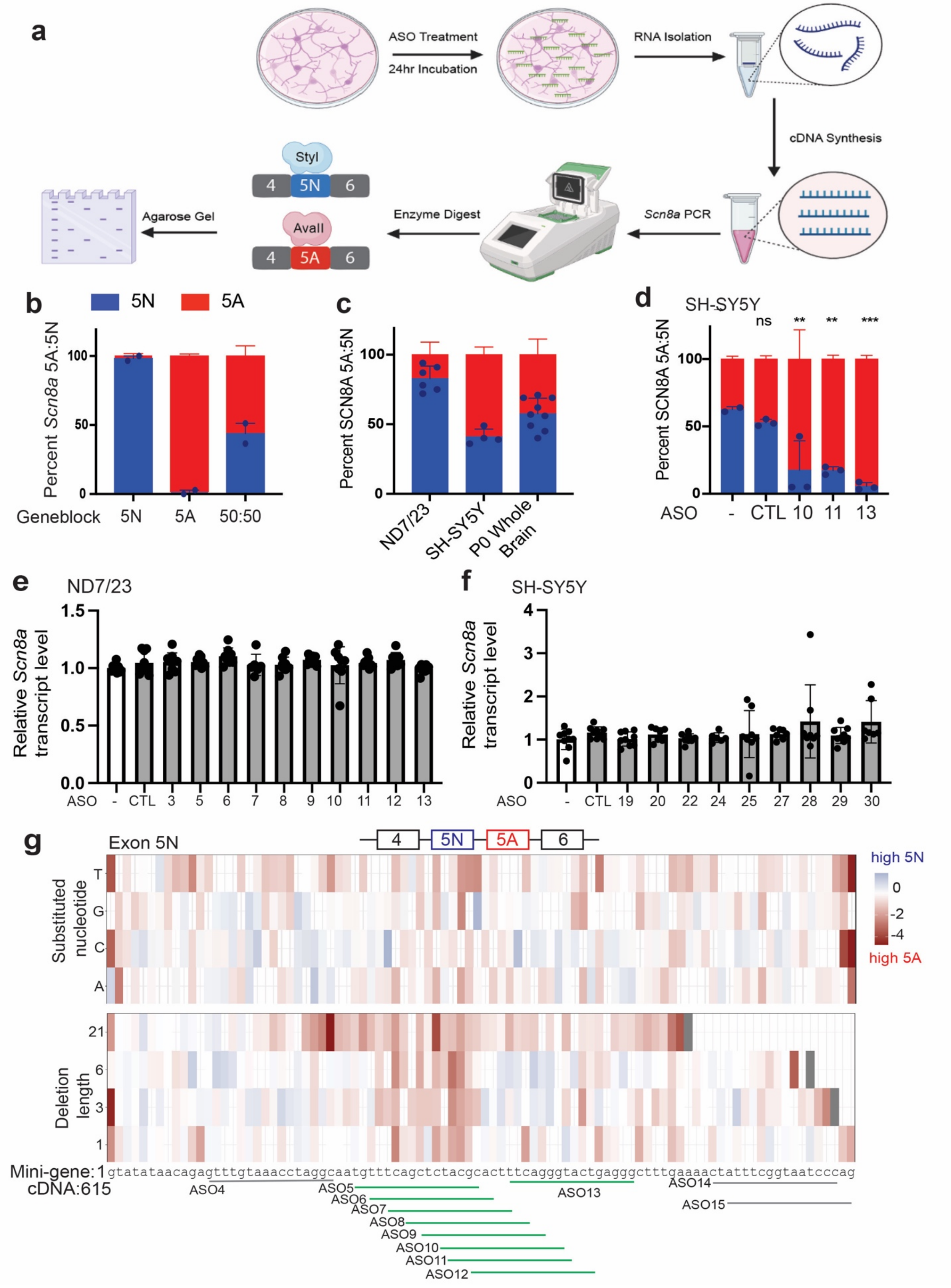

a
ASO Treatment
24hr Incubation
RNA Isolation
cDNA Synthesis
Scn8a PCR
Enzyme Digest
Agarose Gel
StyI
4
5N
6
AvaII
4
5A
6
b
5N
5A
Percent Scn8a 5A:5N
Geneblock
5N
5A
50:50
c
Percent SCN8A 5A:5N
ND7/23
SH-SY5Y
P0 Whole Brain
d
SH-SY5Y
ns
**
**
***
Percent SCN8A 5A:5N
ASO
-
CTL
10
11
13
e
ND7/23
Relative Scn8a transcript level
ASO
-
CTL
3
5
6
7
8
9
10
11
12
13
f
SH-SY5Y
Relative Scn8a transcript level
ASO
-
CTL
19
20
22
24
25
27
28
29
30
g
Exon 5N
4
5N
5A
6
high 5N
high 5A
Substituted nucleotide
T
G
C
A
Deletion length
21
6
3
1
Mini-gene:1
cDNA:615
gtatataacagagtttgtaaacctaggcaatgtttcagctctacgcactttcagggtactgagggctttgaaaactatttcggtaatcccag
ASO4
ASO5
ASO6
ASO7
ASO8
ASO9
ASO10
ASO11
ASO12
ASO13
ASO14
ASO15

**Figure S4: *Scn8a* splicing assay validation in vitro. a**) Schematic detailing *Scn8a* 5A:5N Assay. RNA is extracted from ASO-treated cells, reverse-transcribed into cDNA, which undergoes PCR targeting exon 4-6 of *Scn8a*. PCR product is then digested targeting either the 5A (AvaII) or 5N (StyI) isoform and ran on a TBE-PAGE gel to quantify 5A:5N inclusion. **b**) Percent *Scn8a* 5A:5N expression of differing ratios of exclusive *Scn8a* 5A and 5N geneblocks quantified using the 5A:5N ratio assay (n= 2 per group). **c**) Quantification of percent *Scn8a* 5A:5N expression naturally occurring in ND7/23, SH-5Y5Y cells and cortical neurons cultured for 7 days in vitro isolated from P0 mouse brains (ND7/23 n = 6, SH-SY5Y n = 4, P0 whole brain culture n = 9). **d**) Percent SCN8A 5A:5N expression of 1000 nM gymnotic free uptake ASOS in SH-SY5Y cells (n = 3 biological replicates). **e-f**) qPCR of *Scn8a* total transcript levels of 500 nM lipofectamine transfected ASOs in ND7/23 and SH-SY5Y cells at least n=6 biological replicates per condition. qPCR results plotted as mean ± SD relative to untreated SH-SY5Y cells. Ten ASOs were screened. (**g**) Effect on exon 5N enrichment score of individual mutations and deletions of 1, 3, 6 or 21 bp based on SCN8A minigene library, aligned with 5N targeting ASOs. ASOs that significantly reduce 5N expression via gymnotic uptake in primary cortical neurons are in green. TBE PAGE quantification is plotted as mean ± SD for all graphs. Statistics - one-way ANOVA with Dunnett's multiple comparison test to untreated control; ns $p>0.05$, *$p<0.05$. ** $p<0.01$, *** $p<0.001$, **** $p<0.0001$

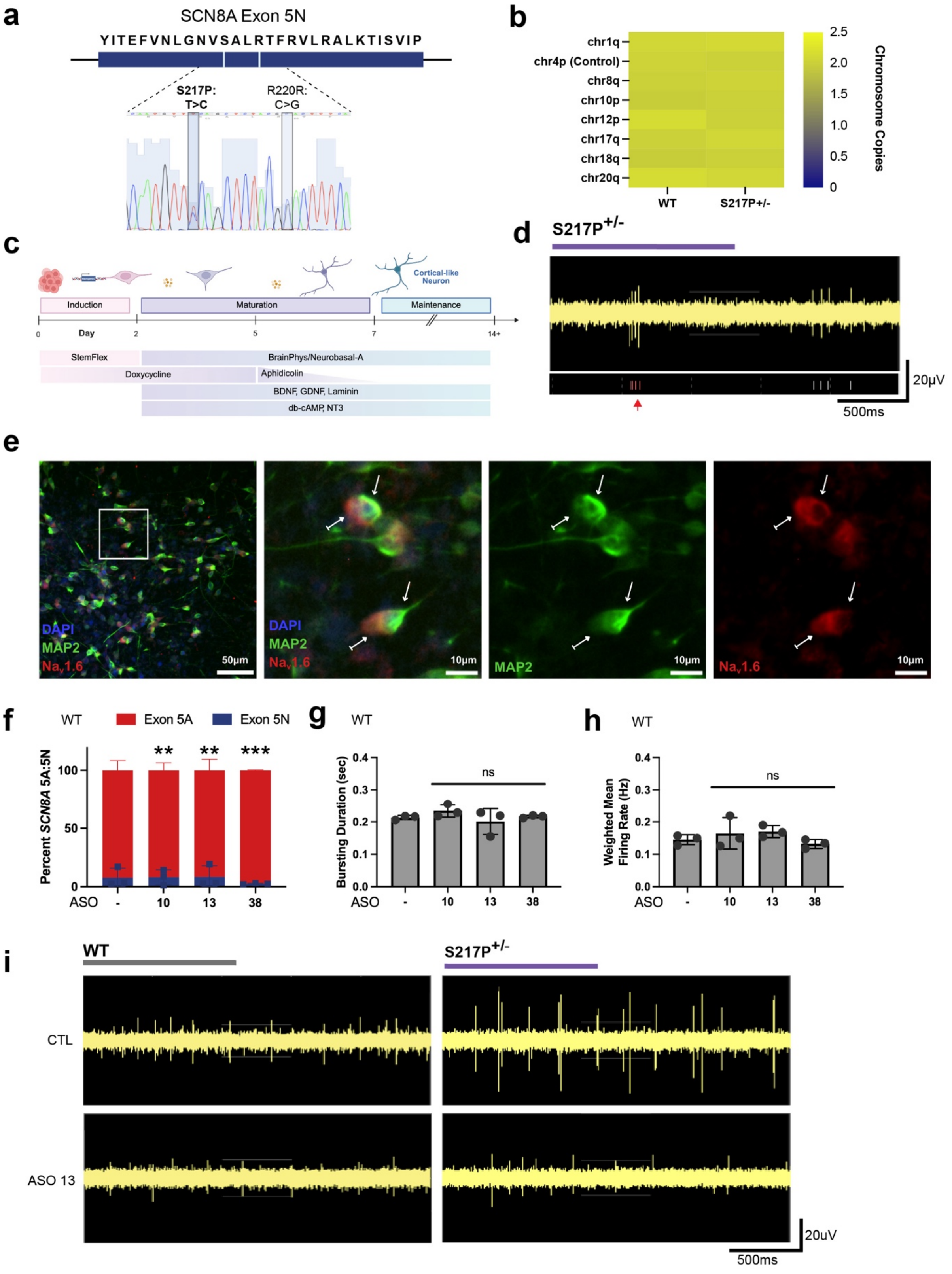


**Figure S5: S217P$^{+/-}$ iPSCs and iPSC-derived induced neurons express standard developmental markers.** **a**, Sanger sequencing of pS217P$^{+/-}$ iPSCs indicates two single nucleotide polymorphisms in SCN8A Exon 5N. Missense T>C at serine 217, and silent C>G at arginine 220. **b**, Representative heat map of chromosomal copy numbers in S217P$^{+/-}$ genome, evaluated by qPCR. Diploid values are indicated in green, no duplications (yellow) or deletions (blue) were detected. **c** Schematic of iPSC-derived neuron induction, differentiation, and maturation. **d**, Representative MEA single electrode raw spikes and associated raster plot from S217P$^{+/-}$ iPSC-neurons indicating a bursting event (red spikes and arrow). **e**, Immunofluorescence of S217P$^{+/-}$ iPSC-neurons at DIV14, stained for the neuronal marker MAP2 (green), SCN8A (red), with DAPI as nuclear reference (blue). The polar distribution of SCN8A is indicated by blunt-end arrows near the axon hillock. MAP2 signal is cytosolic, microtubule densities are indicated by round-end arrows. **f**, Relative percent SCN8A 5A:5N expression in DIV18 WT iPSC-neurons 5 days after 500nM ASO treatment by RNAiMAX transfection. **g-h,** Neuronal network activity of WT iPSC-neuron cultures at DIV18 after 5 days treatment with 500nM ASO, (**g**) bursting duration and (**h**) weighted mean firing rate. **i**, Representative MEA single electrode raw spikes from iPSC-neurons at DIV18 3 days after 500nM ASO treatment. For all data, each point is the mean of at least three technical replicates per biological sample, with error bars plotted as ±SEM. Student's t-test was performed when data were normally distributed (Kolmogorov–Smirnov normality test), and Mann–Whitney U was performed when data were not normally distributed all relative to untreated control; ns p>0.05, *p ≤ 0.05, **p ≤ 0.01, ***p ≤ 0.001, ****p≤0.0001.

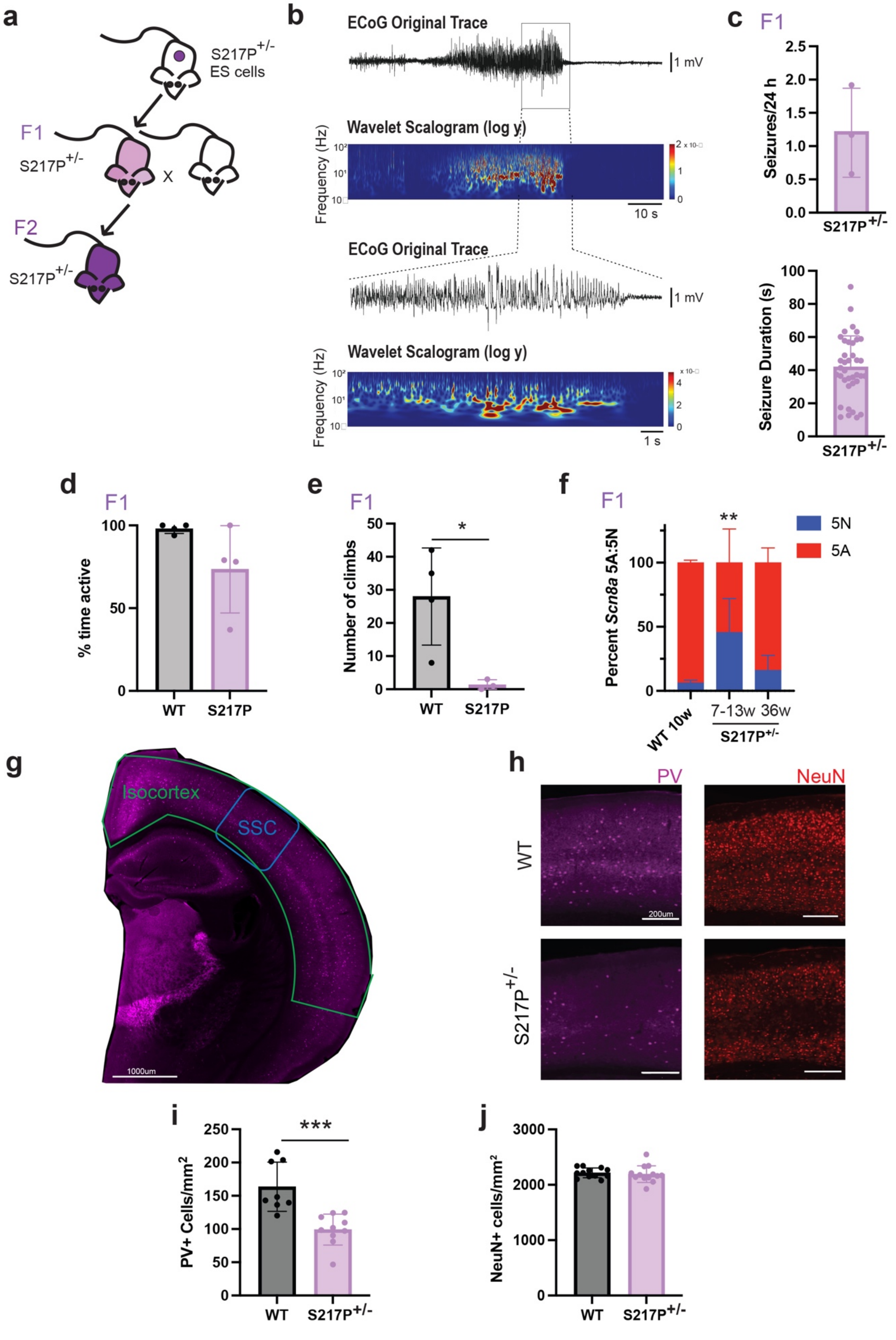

a
S217P+/- ES cells
F1
S217P+/-
X
F2
S217P+/-
b
ECoG Original Trace
1 mV
Wavelet Scalogram (log y)
Frequency (Hz)
10 s
ECoG Original Trace
1 mV
Wavelet Scalogram (log y)
Frequency (Hz)
1 s
c
F1
Seizures/24 h
S217P+/-
Seizure Duration (s)
S217P+/-
d
F1
% time active
WT
S217P
e
F1
Number of climbs
*
WT
S217P
f
F1
Percent Scn8a 5A:5N
**
5N
5A
WT 10w
7-13w
36w
S217P+/-
g
Isocortex
SSC
1000um
h
PV
NeuN
WT
S217P+/-
200um
i
PV+ Cells/mm2
***
WT
S217P+/-
j
NeuN+ cells/mm2
WT
S217P+/-

**Figure S6: Characterization of SCN8A S217P$^{+/-}$ F1 generation.** **a**) Cartoon schematic of the generation of F1 and F2 S217P$^{+/-}$ mice from embryonic cell transfer into wildtype mice. **b-c**) ECoG recordings were conducted on 10-13 week old S217P$^{+/-}$ male mice. Representative trace shows a seizure occurring (**b**). Quantification of seizure occurrence and duration (**c**) (n=3). **d-e**) Quantification of activity and ambulation of adult male wild-type and S217P$^{+/-}$ mice from video recording, n=8 (WT = 4, S217P$^{+/-}$ = 4). **f**) Quantification of 5A:5N inclusion in adult WT (10w) and S217P$^{+/-}$ (7-13w) and aged S217P$^{+/-}$ (36w) mice, n= 17 (WT = 6, S217P$^{+/-}$ 7-13w = 3, S217P$^{+/-}$ 7-13w = 8). **g-j**) Immunostaining (somatosensory cortex representative images – blue outline) and quantification (Isocortex – green outline) for Parvalbumin interneurons (**h-i**) and NeuN (**h-j**) of 10-13 week old S217P$^{+/-}$ mutants and WT C57BL/6 males n= 3 animals per group, 2-4 slices/animal. Statistics – (**d-f**) Unpaired T test, (**h,j**) Linear mixed modeling, comparing genotype, slices are technical replicates to biological animal; ns P>0.05, * P≤0.05, ** P ≤0.01, *** P≤0.001, ****P≤0.0001.

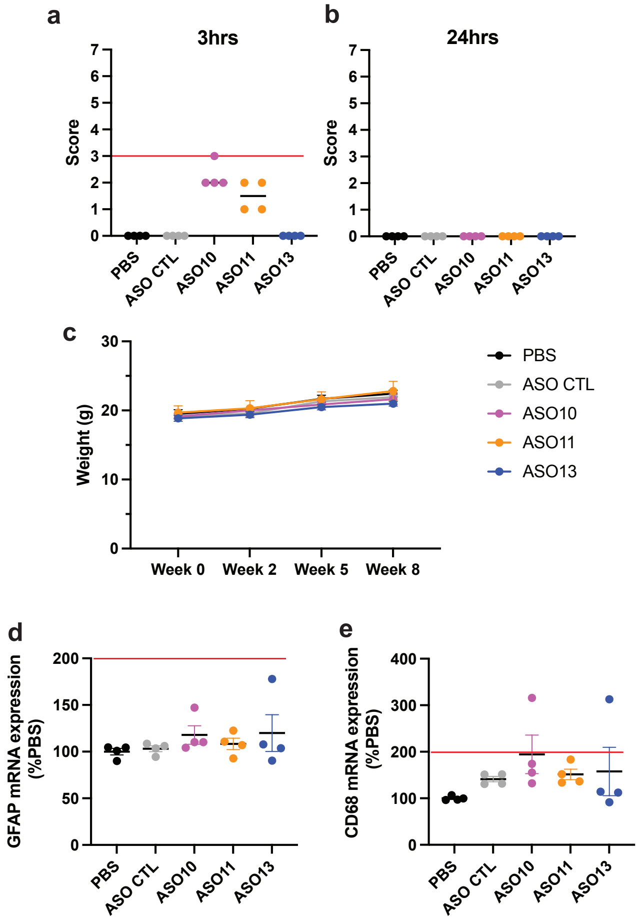


**Figure S7: 5N Targeting ASOs in vivo are non-toxic.** Quantification of acute neuronal activation response (**a**) 3hrs and (**b**) 24hrs after injection of 700ug of ASO ICV in 8-12 week adult mice n=4 mice per group. **c**) Body weight gain over 8 weeks. mRNA expression of (**d**) *Gfap* or (**e**) CD68 in the cortex after 8 weeks of ICV 700ug dose administration of ASO. Red lines represent the maximum limit for toxicity.

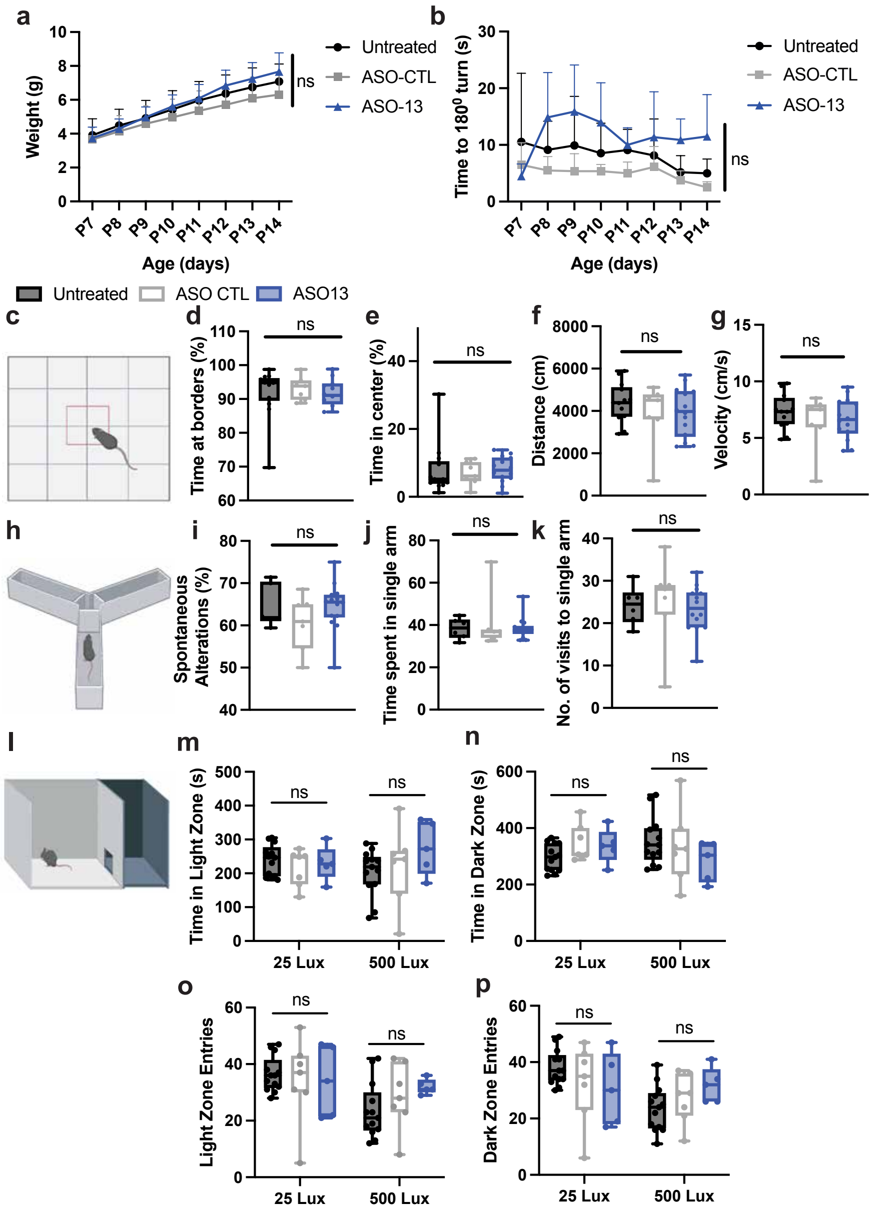


**Figure S8: 5N Targeting ASOs in wild-type animals.** WT mice were treated at P1 with 60ug ASO via ICV injection. (**a-b**) Pups were weighed and tested for negative geotaxis daily from P7 to P14, n= 24 (Untreated = 8, ASO-CTL = 8, ASO-13 = 8). (**c-g**) Six-week old treated animals were observed in the open field for 10 minutes, n= 34 (Untreated = 13, ASO-CTL = 7, ASO-13 = 14). (**h-k**) Six-week old treated animals were observed in the y-maze for 10 minutes, n= 27 (Untreated = 6, ASO-CTL = 7, ASO-13 = 14). (**l-p**) Ten-week old mice were trialed in the light dark box at ambient light (25Lux) on day 1 followed by bright light (500Lux) on day 2, n= 24 (Untreated = 13, ASO-CTL = 7, ASO-13 = 5). Statistics – (**a-b**, **m-p**) two-way ANOVA with Tukey's multiple comparison test comparing all groups. (**d-g**, **i-k**) one-way ANOVA with Dunnett's multiple comparison test to untreated control; ns p>0.05.

**Figure S9: ASO13-tre**

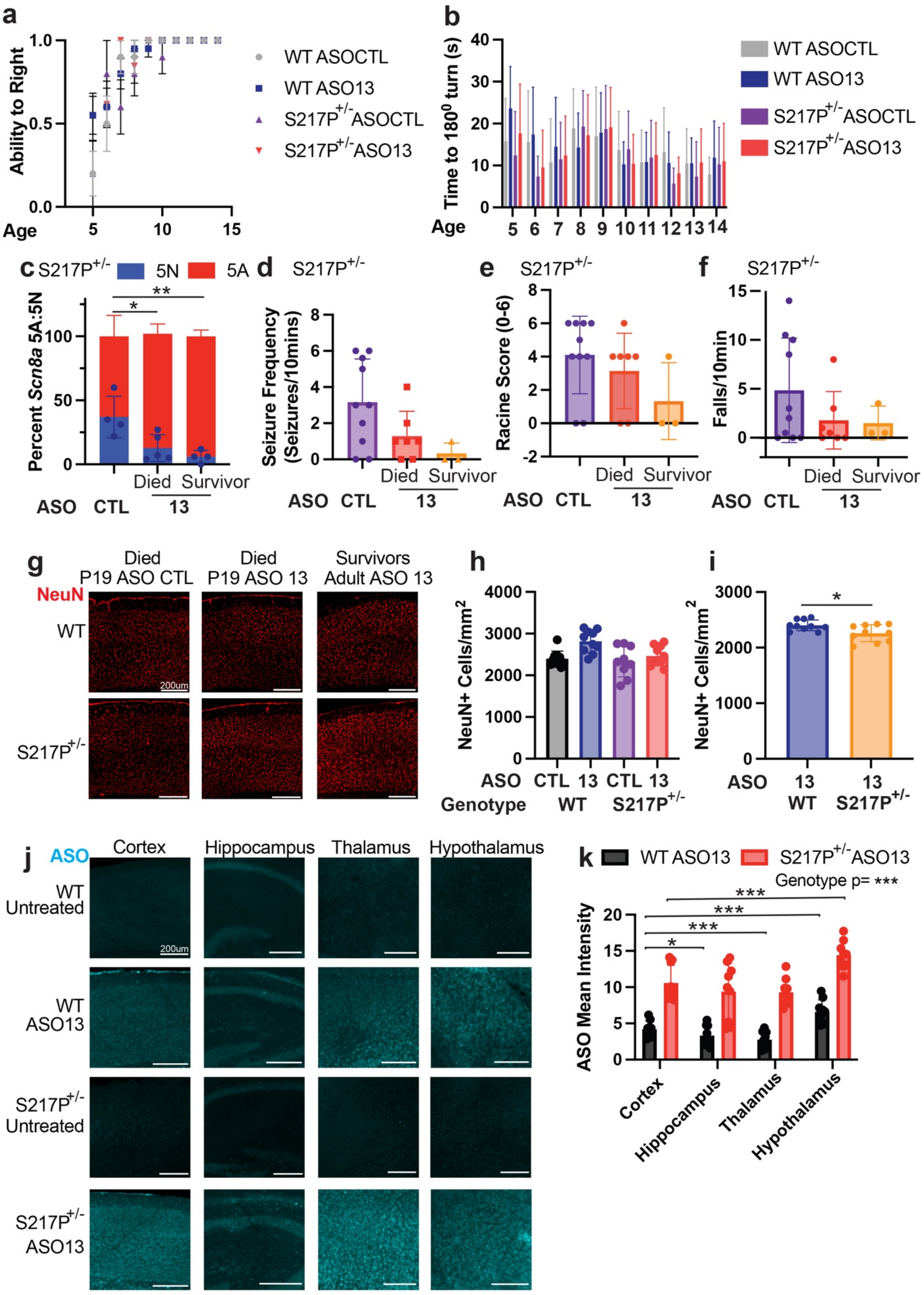
a
Ability to Right
Age
WT ASOCTL
WT ASO13
S217P+/- ASOCTL
S217P+/- ASO13
b
Time to 180⁰ turn (s)
Age
c S217P+/-
5N
5A
Percent Scn8a 5A:5N
Died
Survivor
ASO CTL 13
d S217P+/-
Seizure Frequency (Seizures/10mins)
e S217P+/-
Racine Score (0-6)
f S217P+/-
Falls/10min
g
Died P19 ASO CTL
Died P19 ASO 13
Survivors Adult ASO 13
NeuN
WT
S217P+/-
200um
h
NeuN+ Cells/mm2
ASO CTL 13 CTL 13
Genotype WT S217P+/-
i
ASO 13 13
WT S217P+/-
j
ASO
Cortex
Hippocampus
Thalamus
Hypothalamus
WT Untreated
WT ASO13
S217P+/- Untreated
S217P+/- ASO13
k
WT ASO13
S217P+/- ASO13
Genotype p= ***
ASO Mean Intensity

**ated S217P$^{+/-}$ surviving mice have reduced 5N inclusion, seizures, and impaired movement phenotype compared to ASO-CTL and ASO13 treated S217P$^{+/-}$ mice that died.** Mice were treated at P1 with 60 µg ASO via ICV injection. **a-b**) Pups were tested for righting ability (**a**) and negative geotaxis (**b**), daily from P5-P14, n = 52 (WT ASO-CTL = 10, WT ASO-13 = 20, S217P$^{+/-}$ ASO-CTL = 9, S217P$^{+/-}$ ASO-13 = 13). **c**) *Scn8a* 5A:5N quantification of S217P$^{+/-}$ ASO-CTL and ASO-13 treated animals at death (P16-P23) and endpoint (10-13w), n= 13 (ASO-CTL = 4, ASO13 Died = 5, ASO-13 Survivor = 4). **d-f**) Quantification of P16 video analysis for seizure frequency (**d**), Racine scoring (**e**), and falls (**f**), in S217P$^{+/-}$ treated animals, n = 20 (ASO-CTL = 10, ASO13 Died = 7, ASO13 Survivor = 3). (**g**) Representative images of NeuN positive cells, red, across groups. Scale bar, 200um. (**h**) Quantification of NeuN positive cells/mm$^2$ in ASO treated WT and S217P$^{+/-}$ animals that were sacrificed at mean of P19, *n* = 18 animals (WT ASO-CTL = 5, WT ASO-13 = 5, S217P$^{+/-}$ ASO-CTL = 4, S217P$^{+/-}$ ASO-13 = 4, 1-3 slices/animal). (**i**) Quantification of NeuN positive cells/mm$^2$ in ASO treated WT and S217P$^{+/-}$ adult surviving mice (10-13 weeks old), *n* = 7 animals (WT ASO-13 = 4, S217P$^{+/-}$ ASO-13 survivors = 3, 1-4 slices/animal). (**j**) Representative images of ASO13 staining, cyan, across groups. Scale bar, 200um. (**k**) Quantification of ASO13 intensity ASO13 treated WT and S217P$^{+/-}$ animals that were sacrificed at mean of P19, *n* = 6 animals (WT ASO-13 = 3, S217P$^{+/-}$ ASO-13 = 3, 3 slices/animal).Statistics – (**c**) one-way ANOVA with Dunnett's multiple comparison test to S217P$^{+/-}$ ASOCTL, (**h-j**) Linear mixed modeling, comparing genotype and treatment, slices are technical replicates to biological animal, (j-k) Linear mixed modeling, comparing genotype and region to cortex, slices are technical replicates to biological animal, ns $P>0.05$, * $P\leq0.05$, ** $P \leq0.01$, *** $P\leq0.001$, ****$P\leq0.0001$.

**Figure S10: Percent identity matrices**

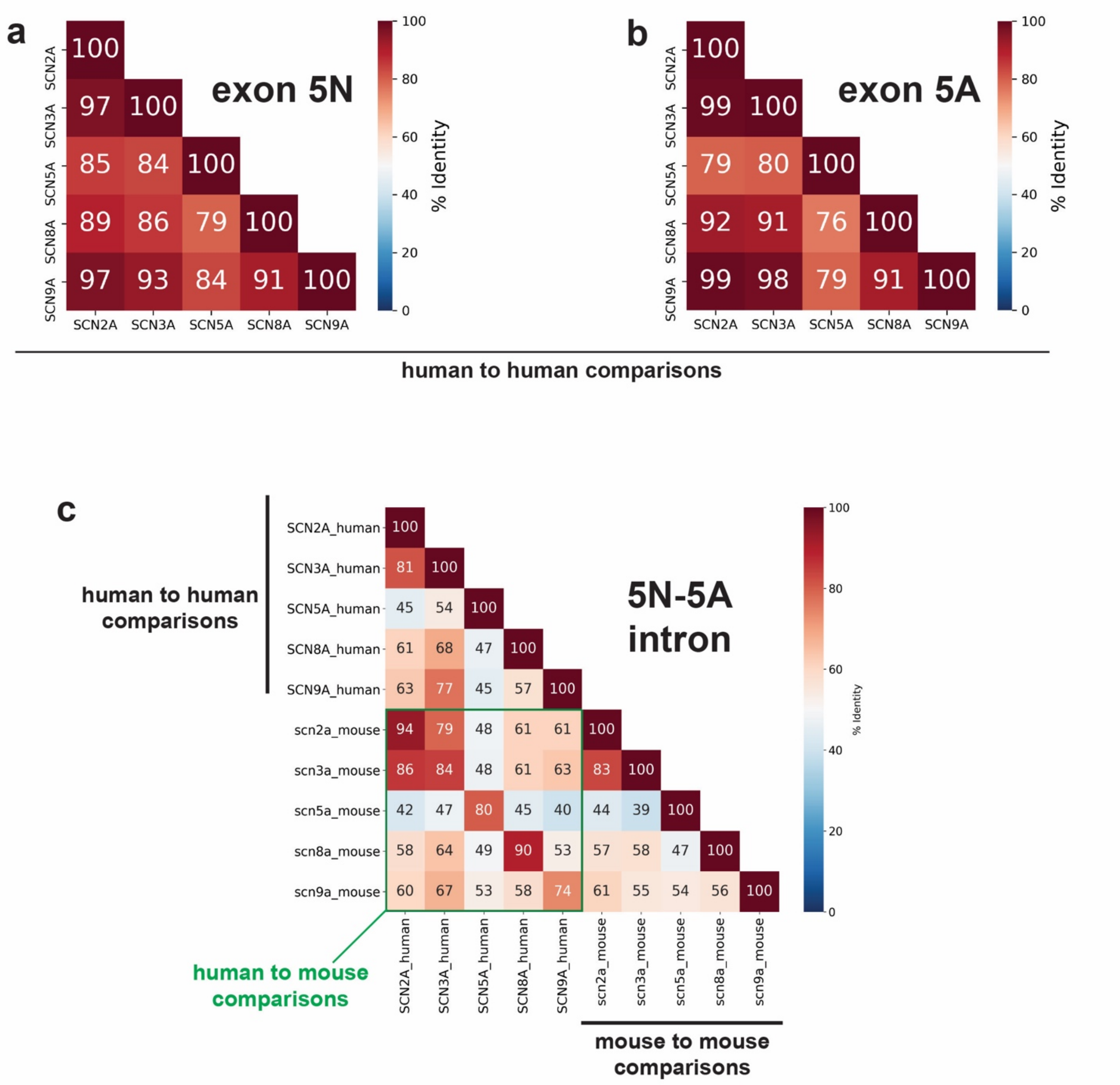


for SCN8A exon 5N **a)**, 5A **b)** from human, and the intron present between 5N and 5A **c)** from human and mouse paralogs. Values represent the % nucleotide sequence identity between sodium channel paralogs as determined by a multiple sequence alignment performed using CLUSTAL Omega. In c) the upper corner depicts comparisons between human paralogs only, the bottom right depicts mouse paralog comparisons only and the green boxed region depicts comparisons between the human and mouse paralogs.
